# ALS and FTD-associated missense mutations in TBK1 differentially disrupt mitophagy

**DOI:** 10.1101/2021.04.12.439050

**Authors:** Olivia Harding, Chantell S. Evans, Junqiang Ye, Jonah Cheung, Tom Maniatis, Erika L.F. Holzbaur

**Affiliations:** Department of Physiology, Perelman School of Medicine, University of Pennsylvania, Philadelphia, PA 19104; Aligning Science Across Parkinson’s (ASAP) Collaborative Research Network, Chevy Chase, MD; Department of Biochemistry and Molecular Biophysics, Columbia University Vagelos College of Physicians and Surgeons, New York, NY 10032; Zuckerman Mind Brain and Behavior Institute, Columbia University, New York, NY 10027; Special Projects Group, New York Structural Biology Center, New York, NY 10027; New York Genome Center, New York, NY 10013

**Keywords:** mitophagy, TBK1, OPTN, Parkin, neurodegeneration

## Abstract

TANK-binding kinase 1 (TBK1) is a multi-functional kinase with an essential role in mitophagy, the selective clearance of damaged mitochondria. More than 90 distinct mutations in TBK1 are linked to amyotrophic lateral sclerosis (ALS) and fronto-temporal dementia (FTD), including missense mutations that disrupt the ability of TBK1 to dimerize, associate with the mitophagy receptor optineurin (OPTN), auto-activate, or catalyze phosphorylation. We investigated how ALS-associated mutations in TBK1 affect Parkin-dependent mitophagy using imaging to dissect the molecular mechanisms involved in clearing damaged mitochondria. Some mutations cause severe dysregulation of the pathway, while others induce limited disruption. Mutations that abolish either TBK1 dimerization or kinase activity were insufficient to fully inhibit mitophagy, while mutations that reduced both dimerization and kinase activity were more disruptive. Ultimately, both TBK1 recruitment and OPTN phosphorylation at S177 are necessary for engulfment of damaged mitochondra by autophagosomal membranes. Surprisingly, we find that ULK1 activity contributes to the phosphorylation of OPTN in the presense of either WT- or kinase inactive TBK1. In primary neurons, TBK1 mutants induce mitochondrial stress under basal conditions; network stress is exacerbated with further mitochondrial insult. Our study further refines the model for TBK1 function in mitophagy, demonstrating that some ALS-linked mutations likely contribute to disease pathogenesis by inducing mitochondrial stress or inhibiting mitophagic flux. Other TBK1 mutations exhibited much less impact on mitophagy in our assays, suggesting that cell-type specific effects, cumulative damage, or alternative TBK1-dependent pathways such as innate immunity and inflammation also factor into the development of ALS in affected individuals.

**SIGNIFICANCE STATEMENT:** Missense mutations in TANK-binding kinase 1 (TBK1) have various biophysical and biochemical effects on the molecule, and are associated with the neurodegenerative diseases amyotrophic lateral sclerosis (ALS) and fronto-temporal dementia (FTD). TBK1 plays an essential role in clearing damaged mitochondria. Here, we investigate the impact of 10 ALS-linked TBK1 mutations on the critical early stage of mitophagy. We find that both TBK1 recruitment and kinase activity contribute to the clearance of the damaged mitochondria. Furthermore, in neurons, expression of TBK1 mutants alone affects mitochondrial network health. Our investigation utilizes disease-linked mutations to further refine the current model of mitophagy, identifying crosstalk between the regulatory kinases TBK1 and ULK1, and providing new insights into the roles of TBK1 in neurodegenerative pathogenesis.

## INTRODUCTION

TNF receptor-associated family member-associated NF-κB activator (TANK)-binding kinase 1 (TBK1) plays a critical role in several cellular pathways implicated in the neurodegenerative disease amyotrophic lateral sclerosis (ALS), including selective clearance of mitochondria and regulation of inflammation. More than 90 mutations in TBK1 have been linked to ALS, including several mutations identified in patients with the co-occurring degenerative disease, fronto-temporal dementia (ALS-FTD) (1, 2). Some TBK1 mutations are classified as loss of function variants while others are missense mutations with unclear contributions to disease pathogenesis (1, 3–6). The latter category includes mutations shown to disrupt the ability of TBK1 to dimerize, associate with the mitophagy receptor optineurin (OPTN), auto-activate, or catalyze phosphorylation (7–9). Given the importance of TBK1 in mitophagy (10), and the necessity of mitochondrial quality control to the maintenance of neuronal homeostasis (11, 12), functional analysis of ALS-associated missense mutations in TBK1 is necessary to determine the impact of mutant TBK1 in the neurodegeneration characteristic of ALS.

TBK1 has three primary domains, 1) a kinase domain, 2) a ubiquitin-like domain, and 3) a scaffold dimerization domain, which are followed by a flexible C-terminus domain (CTD) (Figure 1A) (13–15). Two TBK1 monomers dimerize along their scaffold dimerization domains, while kinase activity is activated via auto-phosphorylation of the critical serine residue 172 (S172) within the activation loop of the kinase domain (14). Due to the conformation of the TBK1 dimer, it is unlikely that the monomers within a dimer can self-activate, so multimer formation is thought to be required for trans-auto-phosphorylation and kinase activation (13, 14). TBK1 multimerization may be promoted by association of TBK1 via its CTD with adaptor proteins including OPTN, TANK, Sintbad, and NAK-associated protein 1 (NAP1) (7, 16, 17). ALS-linked missense mutations are distributed throughout the protein, with some mutations disrupting dimerization, kinase activity, or both, and others disrupting the association of TBK1 with adaptors, potentially inhibiting TBK1 multimerization and activation (Figure 1B) (3, 6–8).

**Figure 1.**
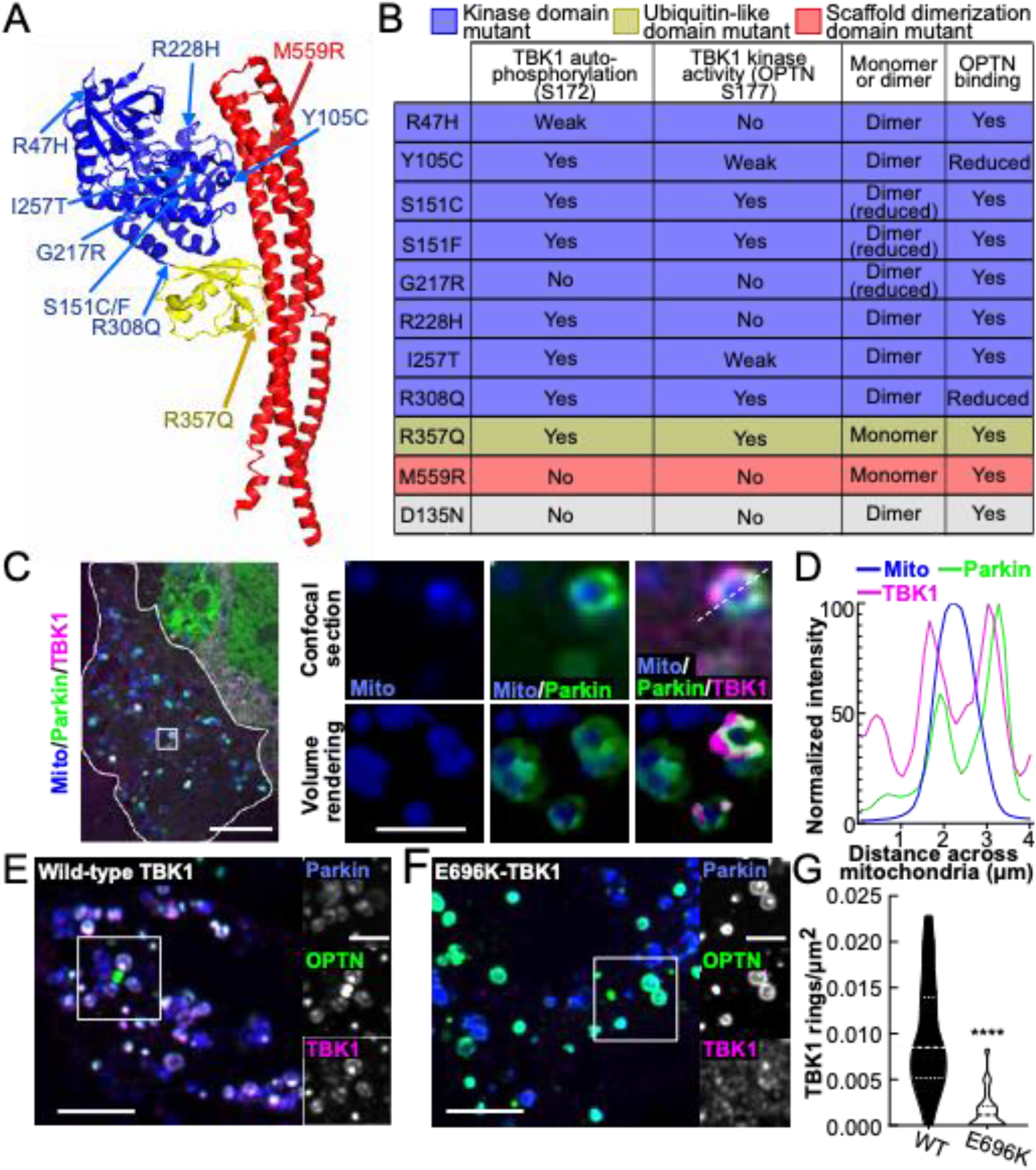
ALS-linked TBK1 mutations are found throughout the molecule and induce biochemical, biophysical, and cellular deficits. A. Protein databank structure for TANK-Binding Kinase 1 (TBK1) (PDB 4IWO) (13). Domains are designated by color coding: kinase domain residues 1-308 (blue), ubiquitin-like domain residues 309-387 (yellow), and scaffolding dimerization domain residues 388-657 (red). ALS-linked mutations are indicated by arrows and labels of their respective colors. Notably, some mutations likely disrupt the structure of TBK1, a phenomenon not represented by this model. B. Table summarizing biochemical results for the ALS-linked mutants published by Ye et al (8), and the engineered kinase-inactive D135N-TBK1 (gray). C. Confocal section of a HeLa cell (outlined in white) expressing a mitochondria-localized fluorophore (blue), Parkin (green) and WT-TBK1 (magenta), fixed after treatment with CCCP for 90 min. The inset (white box) and zoom images (right) exhibit rounded mitochondria that have recruited Parkin and TBK1. A volume rendering is also shown (right, bottom row). Scale bars: zoom out, 10 μm; zoom in, 2 μm. D. Relative signal intensities for mitochondria, Parkin, and TBK1 are quantified across the diameter of a damaged mitochondria (white dashed line in C, zoom). E,F. HeLa cells with depleted endogenous TBK1 expressing Parkin, OPTN, and WT- (E) or E696K- (F) TBK1 fixed after treatment with CCCP for 90 min. Inset (white box) and zoom images (left) demonstrate multiple rings with co-localized mitophagy components. Scale bars zoom out, 10 μm; zoom in, 4 μm. G. Quantification of E and F as rings/μm^2^ for each cell. n= 22-25 cells from 3 independent experiments. Dashed line, median; dotted lines, 25^th^ and 75^th^ quartiles. **** *p* < 0.0001 by student’s unpaired t test. Images E and F shown here are insets; for representative images of whole fields, see Supplemental Figure 1F.

TBK1 kinase activity is an essential regulator of mitophagy, a stepwise pathway for clearance of damaged mitochondria (10, 18). Mitophagy is triggered by loss of mitochondrial membrane potential, leading to the stabilization of PTEN-induced putative kinase 1 (PINK1) on the outer mitochondrial membrane (OMM) (19) where it phosphorylates ubiquitin (20). Phosphorylated ubiquitin recruits the E3 ubiquitin ligase, Parkin (20, 21), which is activated by PINK1 phosphorylation and then ubiquitinates OMM proteins (20, 22–25). These modifications promote proteasomal degradation of mitofusins, preventing the damaged organelle from re-fusing with the healthy mitochondrial network and resulting in a small, rounded mitochondrion (26). Ubiquitination of OMM proteins also promotes recruitment of the mitophagy receptors OPTN, nuclear dot 52 kDa protein (NDP52), Tax1-binding protein 1 (TAX1BP1), next to BRCA gene 1 protein (NBR1), and p62/sequestosome1 (10, 27–30), though OPTN and NDP52 are sufficient and redundant in carrying out mitochondrial clearance in HeLa cells (29). Phosphorylation of OPTN at S177 by TBK1 at the OMM enhances the binding of OPTN to ubiquitin chains (18). OPTN then drives recruitment of the core autophagy machinery, including the unc-51-like autophagy activating kinase (ULK1) complex, to initiate formation of the double membraned phagophore that engulfs the damaged organelle (31–33). In this process microtubule-associated protein 1A/1B-light chain 3 (LC3) is lipidated and subsequently incorporated into the elongating phagophore (10, 27, 34). The LC3-interacting region of OPTN facilitates efficient engulfment by the autophagosome (10), while TBK1-mediated phosphorylation of OPTN enhances the binding of the receptor to LC3 (35). A feed-forward mechanism in which initial LC3-positive membranes recruit more OPTN and NDP52 leads to accelerated mitochondrial engulfment (36). The newly formed compartment fuses with lysosomes to complete degradation of the organelle (30, 37, 38).

We undertook a functional analysis of ALS-associated TBK1 missense mutations that have been characterized by biochemical and biophysical assays but confer unknown effects on the cellular pathways that involve TBK1. We determined the extent of recruitment of TBK1 mutants to depolarized, Parkin-positive mitochondria, the effect of mutant TBK1 expression on OPTN recruitment and phosphorylation, and the resulting downstream engulfment of fragmented mitochondria by LC3-positive autophagosomes. Expression of some ALS-linked mutations profoundly disrupted TBK1 recruitment and activity during mitochondrial clearance, while others only marginally affected the pathway. Neurons expressing TBK1 mutations demonstrated higher baseline levels of mitochondrial stress and an inability to manage induced oxidative damage, both of which may contribute to neurodegeneration. Our data suggest a more nuanced model of TBK1 function, wherein TBK1 phosphorylates OPTN directly, while TBK1 recruitment also facilitates OPTN phosphorylation via an ULK1 dependent pathway. Further, we demonstrate that ALS and ALS-FTD-associated missense mutations in TBK1 can lead to disordered or delayed mitochondrial clearance and a cellular deficiency in mitochondrial homeostasis.

## RESULTS

In order to test whether mutations in TBK1 affect mitophagy, we used a well-characterized assay in HeLa-M cells, in which mitochondria were depolarized with the mitochondrial membrane disrupter, carbonyl-cyanide *m*-chlorophenyl-hydrazone (CCCP), and components of the mitophagy pathway were visualized by fluorescent microcopy (10, 27). We depleted cells of endogenous TBK1 and expressed SNAP- or Halo-tagged TBK1 (Supplemental Figure 1A,B) along with a fluorescently-tagged mitochondrial marker, Parkin, OPTN, or LC3. While some constructs had a lower transfection efficiency as compared to WT-TBK1, most were expressed at similar cellular levels, with the exceptions of S151F and M559R, which exhibited slightly but statistically higher cellular expression (Supplemental Figure 1C-E). Under basal conditions, TBK1 was mostly cytosolic with intermittent puncta that did not associate with Parkin (Supplemental Figure 1F).

With 90 minutes of CCCP treatment, Parkin, OPTN, TBK1, and LC3 assembled in a molecular platform at the OMM that appears as a ring surrounding a rounded mitochondrion in single plane confocal sections (10); in Z-stacks the complete engulfment of the mitochondrion is apparent (Figure 1C,D, Supplemental Figure 2A,B) (30). The time course of ring formation observed in HeLa-M cells overexpressing Parkin is similar to that observed in hippocampal neurons expressing endogenous Parkin (30). E696K is a mutation in TBK1 that was previously shown to inhibit recruitment of TBK1 to mitochondria after depolarization (10). We treated E696K- or WT-TBK1 expressing cells with CCCP and compared the prevalence of TBK1 rings after 90 min (Figure 1E,F Supplemental Figure 2B). E696K-TBK1 expressing cells had significantly fewer rings/μm^2^ than WT-TBK1 expressing cells (Figure 1G), establishing this approach as a quantitative measure of the functional effects of TBK1 on mitophagy. Of note, loss of mitochondrial mass was not observed within this time frame, as lysosomal degradation is not evident until ∼12 hours after induction of mitochondrial damage (26, 27, 29).

### Dimerization mutations do not preclude TBK1 recruitment

TBK1 dimerization is proposed to stabilize the trimodular structure of the molecule and permit efficient activation and kinase activity (13). We asked whether two ALS-associated mutations that prevent dimerization, R357Q and M559R (8), would affect TBK1 recruitment to damaged mitochondria. In basal conditions, SNAP-tagged R357Q- and M559R-TBK1 were cytosolic with intermittent puncta (Supplemental Figure 1F), and their expression did not appreciably affect mitochondrial content. Following CCCP treatment, cells expressing WT-, R357Q-, or M559R-TBK1 exhibited robust Parkin recruitment to rounded mitochondria (Figure 2A, Supplemental Figure 2C). R357Q-TBK1 was recruited to the same extent as WT-TBK1 (Figure 2A,B). Strikingly, despite the higher cellular expression level (Supplemental Figure 1E), no M559R-TBK1 recruitment to mitochondria was evident (Figure 2A,B). Instead, M559R-TBK1 remained largely cytosolic with some apparent aggregate formation, although these aggregates were not associated with mitochondria (Figure 2A).

**Figure 2.**
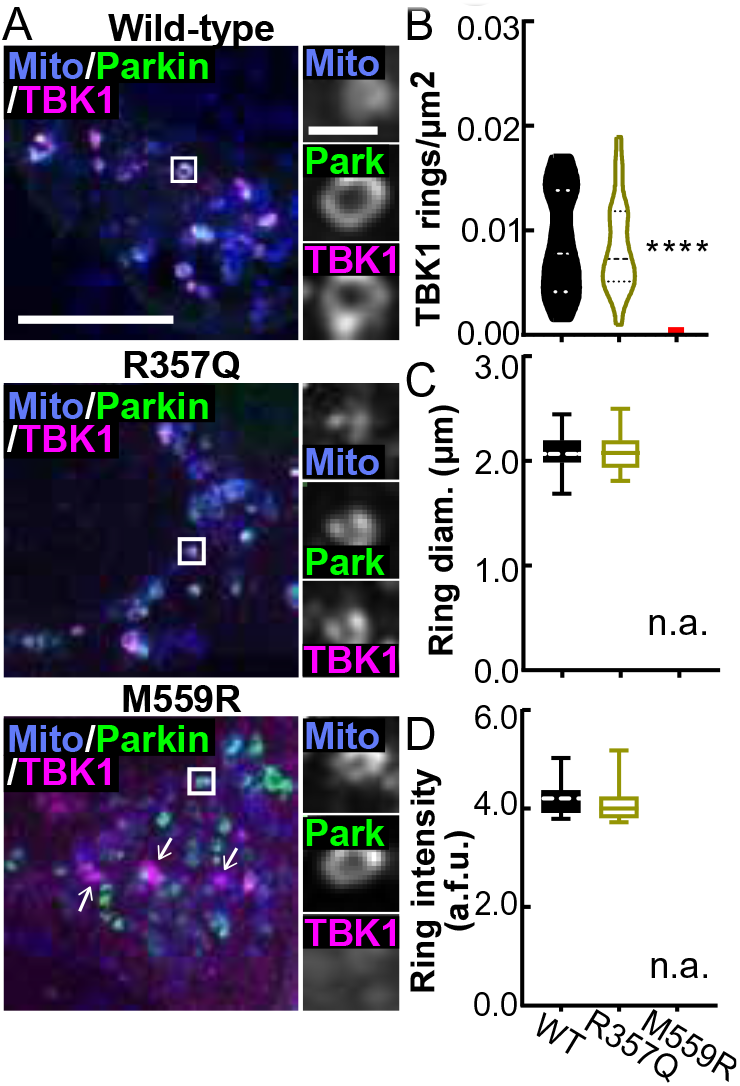
TBK1 mutants that are unable to dimerize are differentially recruited to damaged mitochondria. A. Maximum intensity projection images of fixed HeLa cells depleted of endogenous TBK1 expressing a mitochondria-localized fluorophore (blue), Parkin (green), and WT- (top row), R357Q- (middle row), or M559R- (bottom row) TBK1 (magenta) after 90 min CCCP. There are some aggregates of M559R-TBK1 (arrows) that are not co-incident with mitochondria. Scale bars zoom out, 10 μm; zoom in, 2 μm. Images shown are insets; for representative images of whole fields, see Supplemental Figure 2D. B-D. Quantification of TBK1 rings/μm^2^ (B) ring diameter (C), and ring signal intensity (D). **** *p* < 0.0001 by ordinary one-way ANOVA with Dunnett’s multiple comparisons test. Dashed line, median; dotted lines, 25^th^ and 75^th^ quartiles. No M559R-TBK1 rings were evident, so all data points are zero for rings/μm^2^ (red line) and no data can be displayed for size and intensity. n= 22-26 cells from 3 independent experiments. Data in C-D analyzed by students unpaired t test. Not applicable, n.a. Arbitrary fluorescent units, a.f.u.

We measured the size and intensity of TBK1 rings to assess whether R357Q-TBK1 conferred a structural defect on the ubiquitin-based molecular platform that forms on damaged mitochondria. R357Q-TBK1 rings were the same diameter and average fluorescence intensity as WT-TBK1 rings (Figure 2C,D), indicating that the monomeric property of R357Q-TBK1 does not impair its ability to form the molecular ring structure. Together these observations suggest that the lack of M559R-TBK1 recruitment is not due solely to an inability of the molecule to dimerize.

### Mutations disrupting both dimerization and activation impair recruitment of TBK1 to damaged mitochondria

The M559R mutation in TBK1 also disrupts kinase activation and enzymatic activity (Figure 1B) (8), so we employed our mito-depolarization assay to test other TBK1 missense mutations that exhibit reduced auto-phosphorylation activity to varying degrees: R47H-TBK1, G217R-TBK1, and R228H-TBK1. G217R-TBK1 exhibits reduced dimer formation as well (8, 9). Of the mutants tested, only G217R-TBK1 exhibited deficient recruitment to damaged mitochondria compared WT-TBK1. Expression of G217R-TBK1 resulted in significantly decreased TBK1 ring density, despite clear evidence of mitochondrial fragmentation and Parkin recruitment (Figure 3A,B, Supplemental Figure 3A).

**Figure 3.**
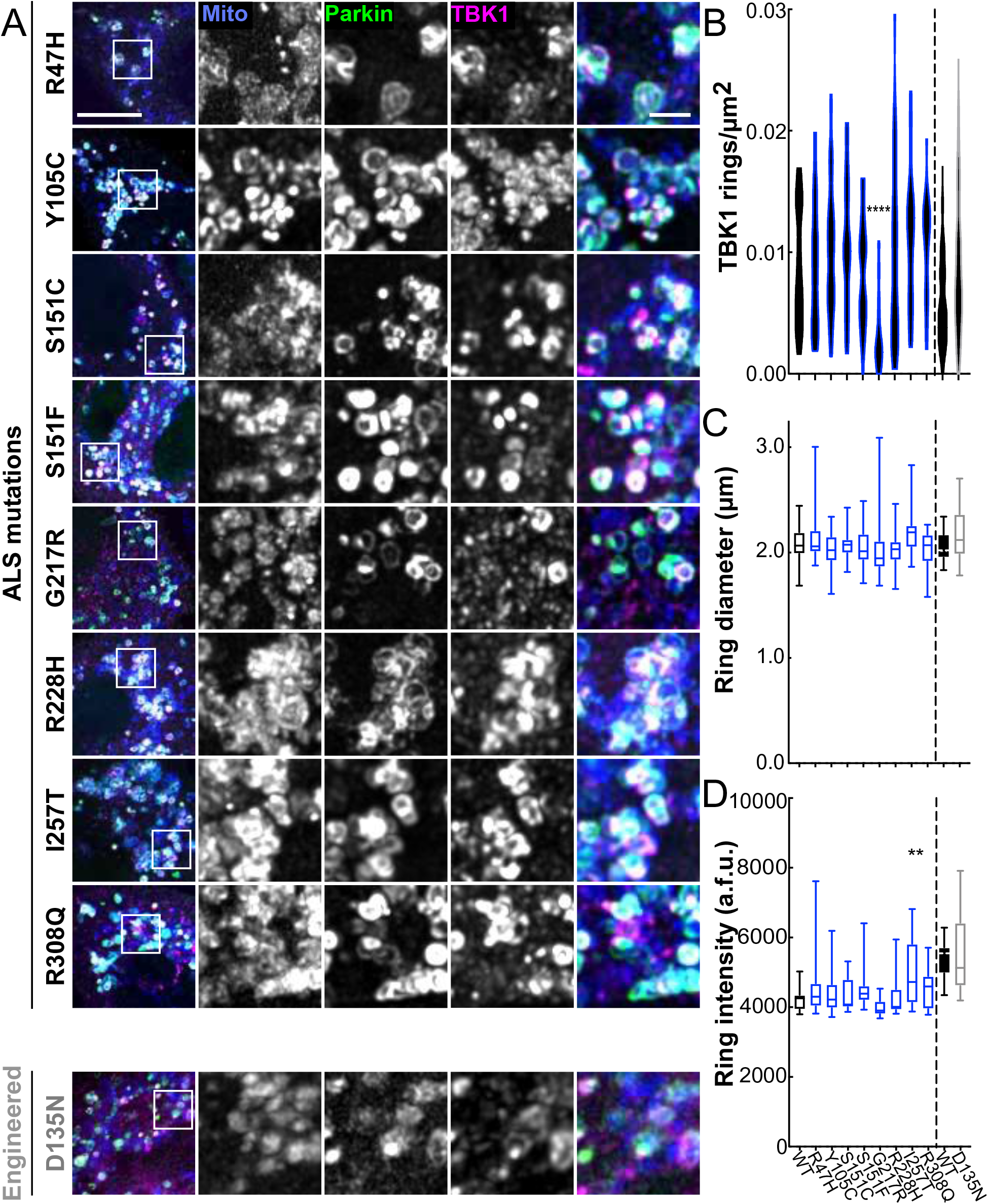
A kinase domain mutation that abolishes the auto-phosphorylation ability of TBK1 results in fewer TBK1 rings. A. Maximum intensity projection images of fixed HeLa cells depleted of endogenous TBK1 expressing fluorescent mitochondrial marker (blue), Parkin (green), and TBK1 variants (magenta) and fixed after treatment with CCCP for 90 min. Images shown are insets; for representative images of whole fields, see Supplemental Figure 3A. B-D. Quantification of TBK1 rings/μm^2^ (B) ring diameter (C), and ring signal intensity (D). WT-TBK1 ring data is transferred from Figure 2 for comparison, indicated by black outline. n= 22-32 cells from at least 3 independent experiments. For ring density (B) dashed line, median; dotted lines, 25^th^ and 75^th^ quartiles. ** p < 0.01, **** p < 0.0001 by ordinary one-way ANOVA with Dunnett’s multiple comparisons test. Arbitrary fluorescent units, a.f.u.

To further delineate the role of kinase activity, we expressed a TBK1 variant with an engineered mutation, D135N, which renders the TBK1 molecule kinase inactive and unable to auto-phosphorylate at S172, but fully able to dimerize (16). In line with previous data on an engineered phospho-deficient S172A-TBK1 mutant (10), D135N-TBK1 was recruited to damaged mitochondria to the same extent as WT-TBK1 (Figure 3A,B). Moreover, the ALS-associated mutations R47H-TBK1 and R228H-TBK1 have weaker auto-phosphorylation activity than WT-TBK1 based on biochemical studies (8), yet they also translocated to damaged mitochondria with the same incidence as WT-TBK1 (Figure 3A,B).

We saw no difference in either ring diameter or intensity across WT-, R47H-, G217R-, and R228H-TBK1 rings (Figure 3C,D), or when comparing WT- and kinase-inactive D135N-TBK1. Only I257T-TBK1, a kinase domain mutant exhibiting WT-TBK1 levels of auto-phosphorylation but weaker kinase activity toward OPTN, formed brighter rings than WT-TBK1. However, the average diameter of the I257T-TBK1 rings were not significantly different from WT-TBK1 rings (Figure 3C,D). Notably, the similarities between WT-TBK1 rings and the poorly recruited G217R-TBK1 rings demonstrate that expression of ALS-associated TBK1 mutants does not disrupt the integrity of TBK1 rings, even if fewer rings form. It is unlikely that mutant TBK1 recruitment is due to dimerization of mutant TBK1 with residual endogenous TBK1, since we measured knockdown levels >70% (Supplemental Figure 1A,B). To further substantiate this claim, we took advantage of the fact that M559R-TBK1 expressing cells exhibit no detectable recruitment of the tagged exogenous construct, and probed CCCP-treated cells expressing M559R-TBK1 with an anti-TBK1 antibody to detect total TBK1. No TBK1 reactivity was detected at damaged mitochondria (Supplemental Figure 3B).

To assess TBK1 recruitment with an alternative method of mitochondrial depolarization, we treated a subset of TBK1 variant-expressing cells with a combination of Antimycin A and Oligomycin A for 90 min (39). A majority of WT-, R357Q-, and D135N-TBK1 expressing cells exhibited TBK1 rings, while significantly less cells expressing G217R-TBK1 (43 ± 12%) or M559R-TBK1 (3.0 ± 3%) exhibited rings (Supplemental Figure 4). Recapitulation of TBK1 recruitment patterns across different depolarizing insults fortifies our finding that R357Q- and D135N-TBK1 are recruited to damaged mitochondria to the same extent as WT-TBK1, while G217R- and M559R-TBK1 are deficient in this step of mitochondrial clearance.

### M559R- and G217R-TBK1 display disrupted recruitment kinetics compared to WT- and R357Q-TBK1

Biochemical analyses indicate that the missense mutations G217R, R357Q, and M559R impair the function of TBK1 (3, 8, 9). Expression of these variants may disrupt recruitment kinetics during individual mitophagic events, as compared to WT-TBK1. We performed live cell microscopy using Halo-tagged TBK1 constructs and tracked single mitophagy events from initial Parkin recruitment to peak TBK1 recruitment in cells expressing similar levels of exogenous TBK1. R357Q-TBK1 exhibited the same kinetics as WT-TBK1 (Figure 4A), reaching maximum intensity as a fully formed ring ∼10-15 min after Parkin reached its half-maximum. Together, the comparative kinetics (Figure 4B) and ring prevalence between WT- and R357Q-TBK1 (Figure 2) suggest that TBK1 dimerization is not required for recruitment and assembly of TBK1 at the depolarized mitochondria.

**Figure 4.**
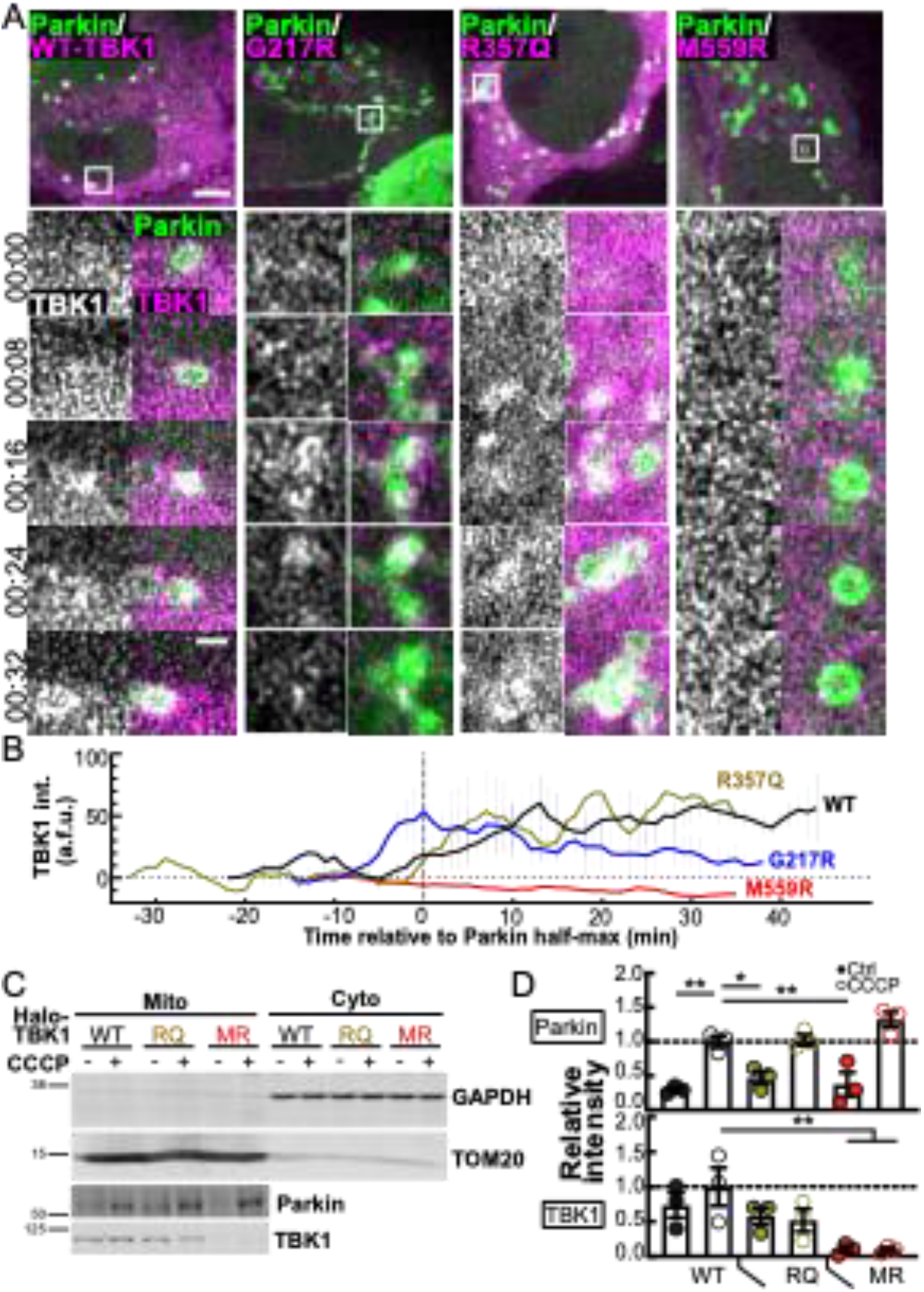
TBK1 variants exhibit differing kinetics and affinities with damaged mitochondria. A. Representative confocal sections of live HeLa cells depleted of endogenous TBK1 and expressing Parkin (green) and TBK1 variants (magenta), treated with CCCP for up to 90 min. White box inset indicates a single representative event tracked over time to measure recruitment of Parkin and TBK1. Stills from timelapse are shown in the panels. Time is indicated as min:sec from initial Parkin recruitment. Scale bars, zoom out, 10 μm; zoom in, 2 μm. B. Background-subtracted signal tracked over time with respect to Parkin half-maximum (0, vertical dotted line). n= 3-6 representative events from at least 3 independent experiments. Error bars indicate SEM. C. A representative Western blot of HEK TBK1−/− cells expressing the respective TBK1 variants, treated with CCCP or vehicle, and enriched for mitochondria (left, “Mito”) or cytosol (right, “Cyto”). Quantification was carried out on mito fractions to compare association of the respective TBK1 variants and Parkin with mitochondria. Numbers to the left of blots indicate kDa based on protein ladder (not shown). D. Quantification of (C) with Parkin (top graph) and TBK1 (bottom graph) bands normalized to TOM20 and compared to average level of WT-TBK1 expressing cells treated with CCCP (dotted line). * p ≤ 0.05, ** p < 0.01 by ordinary one-way ANOVA with Dunnett’s multiple comparisons test. Error bars indicate SEM. n= 3 independent experiments.

G217R-TBK1 translocated to and coalesced at damaged mitochondria in bright but unstable structures at around the same time that Parkin reached its half-maximum (Figure 4A,B). These unstable configurations reached the same raw maximum intensity as WT- and R357Q-TBK1 and occasionally appeared as full rings. Some G217R-TBK1 rings remained intact over the course of our observation (up to 60 min) (Figure 3A), but the majority disappeared within 20 min of reaching peak intensity (Figure 4B). In contrast, over the course of 90 min of mitochondrial damage M559R-TBK1 was never recruited to damaged mitochondria (Figure 4A,B).

The kinetic data (Figure 4A,B) and the results from fixed cells (Figure 2,3) indicate that there is a specific window of time during which TBK1 must be recruited in order to properly form and maintain a stable ring. If such interactions are insufficient, TBK1 molecules disperse from the damaged organelle as we see with G217R-TBK1. The inability of G217R-TBK1 to carry out auto-phosphorylation combined with its reduced dimerization may make a stable interaction with ubiquitinated mitochondria unlikely. Since R357Q-TBK1 can be phosphorylated and activated even as a monomer, it was able to maintain a stable ring structure. M559R-TBK1 is completely monomeric, and it is even less likely to stably interact with the mitochondria, thus there are no detectable rings following mitochondrial damage.

To further probe the integrity of the association of TBK1 with damaged mitochondria, we employed a mitochondrial fractionation assay. We expressed R357Q-, M559R-, or WT-TBK1 in human embryonic kidney (HEK) cells in which both TBK1 alleles had been deleted by CRISPR-Cas9 (HEK TBK1−/−) (8), then enriched mitochondria from the cells after CCCP or vehicle treatment (Figure 4C). CCCP treatment resulted in twice as much Parkin enriched in the mitochondrial fraction of each sample as compared to vehicle treated cells, demonstrating the assay’s sensitivity to Parkin-dependent mitophagy. This Parkin enrichment corroborated our immunofluorescent data in HeLa-M cells as well as previous experiments in HEK cells (38). Expression of R357Q- or M559R-TBK1 did not affect Parkin enrichment after mitochondrial depolarization.

TBK1 is recruited to OPTN puncta after 90 min CCCP treatment in HEK cells, however the effect is less robust than in HeLa cells (Supplemental Figure 4C). In mitochondrial fractions, there was a low affinity association of WT-and R357Q-TBK1 with mitochondria under basal conditions (Figure 4C,D). In contrast, M559R-TBK1 was barely present in the mitochondrial fraction (Figure 4C,D). With CCCP-treatment, we did not detect a significant increase in TBK1 in the mitochondrial fraction with expression of any of the variants, suggesting that TBK1 may be transiently associated with mitochondria even in the absence of induced stress. The absence of M559R-TBK1 under basal or mitochondrial damage conditions substantiates the severe functional defect of M559R-TBK1 seen in HeLa assays.

### OPTN phosphorylation is enhanced by TBK1 recruitment but not fully dependent on TBK1 activity

Activated TBK1 leads to the phosphorylation of OPTN on at least two residues, S177 and S513; these phosphorylation events enhance the binding of OPTN to ubiquitin chains deposited on damaged mitochondria in the early stage of mitophagy (18). However, TBK1 activity is not required for OPTN recruitment to damaged mitochondria, as OPTN was recruited in cells in which TBK1 was either depleted or its kinase activity was inhibited (10) (Supplemental Figure 5A). Further, we found that none of the TBK1 mutants affected OPTN recruitment to damaged mitochondria (Supplemental Figure 5B). In the clearest case, expression of M559R-TBK1 results in a complete absence of TBK1 rings (Figure 2). However, OPTN ring prevalence was not impacted, and TBK1 mutant expression did not cause variations in the size or intensity of OPTN rings, with the exception of a slightly larger diameter of OPTN rings induced by R47H-TBK1 expression (Supplemental Figure 5B,C).

The LC3-interacting region of OPTN is involved in facilitating formation of the LC3-positive autophagosome (35), so both recruitment and phosphorylation of OPTN at S177 may be necessary to complete clearance of damaged mitochondria. We asked whether expression of mutant variants of TBK1 inhibits the phosphorylation of OPTN. To this end, we performed immunofluorescence using an antibody to phospho-S177 OPTN to assess the extent of OPTN modification (Figure 5, Supplemental Figure 6A). WT-TBK1 expressing cells exhibited robust OPTN rings after 90 min of CCCP treatment (Figure 5A). Further, WT-TBK1 colocalized with OPTN rings, and there was a strong phospho-OPTN signal coincident with TBK1-positive OPTN rings (Figure 5A, Supplemental Figure 6A). Over 75% of OPTN rings in each cell were either TBK1-positive, phospho-OPTN-positive, or positive for both; the majority (58 ± 5.8%) of all OPTN rings were coincident with both TBK1 and phospho-OPTN (Figure 5F). R357Q- and D135N-TBK1 expression exhibited similar results (Figure 5F).

**Figure 5.**
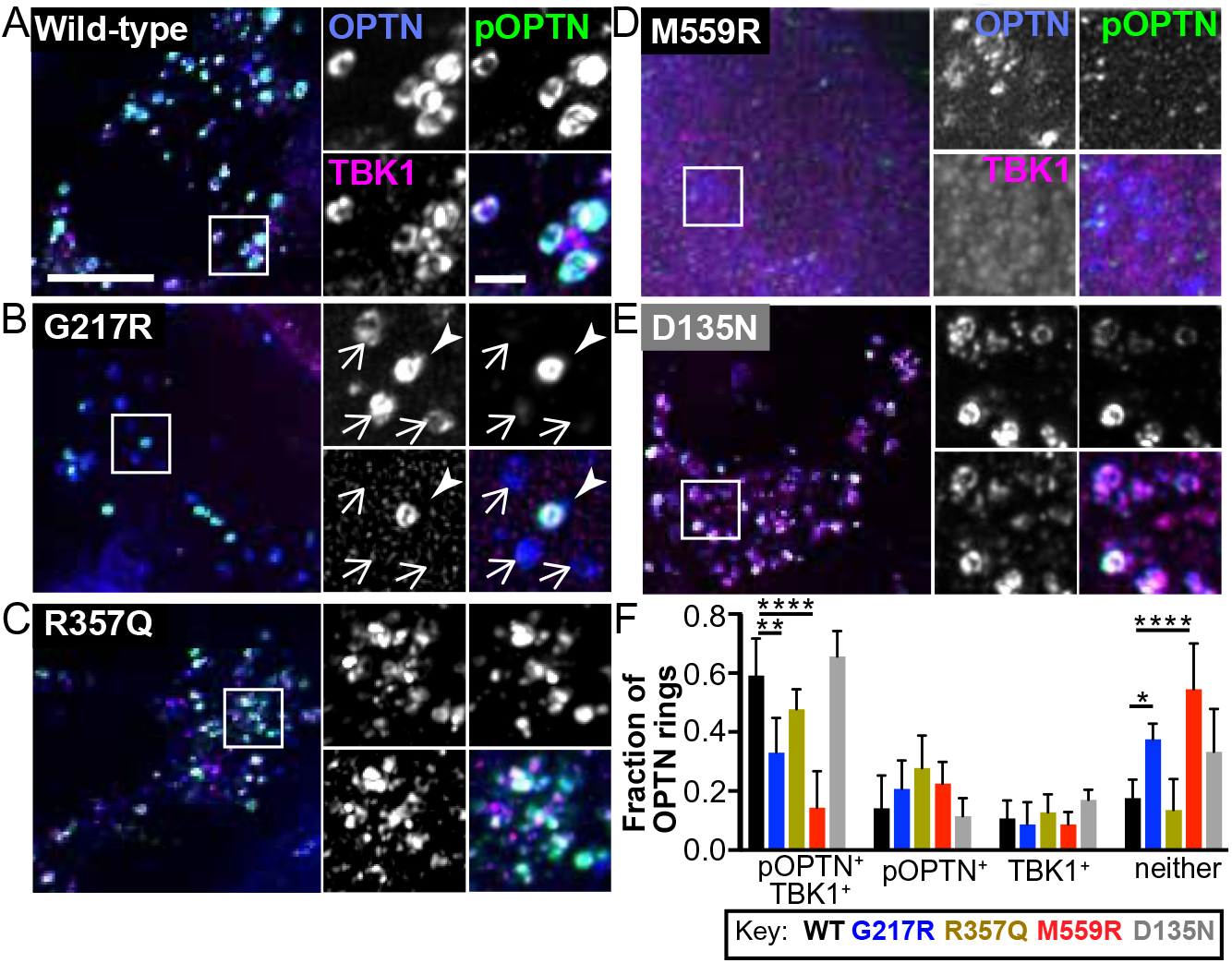
TBK1 mutant expression does not impact OPTN ring incidence. A-E. Maximum intensity projection images of HeLa cells depleted of endogenous TBK1 and expressing Parkin (not tagged), OPTN (blue), and WT- (A), G217R- (B), R357Q- (C), M559R- (D), or D135N- (E) TBK1 variants (magenta), fixed after treatment with CCCP for 90 min. Phospho-S177-OPTN is tagged with an antibody (green). In B, one ring is positive for phospho-OPTN and TBK1 (arrowhead), and the others are negative for both (arrows). Scale bars, zoom out, 10 μm; zoom in, 2 μm. Images shown are insets; for representative images of whole fields, see Supplemental Figure 6A. For each cell the percentage of OPTN rings in each category was calculated, and these results are displayed by bar graph to the right of the respective images. Error bars indicate SD. n= 8-15 cells from at least 3 independent experiments. * p ≤ 0.05, ** p < 0.01, **** p < 0.0001 by ordinary one-way ANOVA with Dunnett’s multiple comparisons test.

G217R- and M559R-TBK1 expressing cells exhibited OPTN rings with lower intensities of TBK1 and phospho-OPTN (Supplemental Figure 6B,C). Consistent with our observations that G217R-TBK1 rings were not as prevalent as WT-TBK1 rings (Figure 3), G217R-TBK1 expressing cells exhibited fewer OPTN rings that were positive for TBK1. However, ∼50% of OPTN rings were positive for phospho-OPTN (Figure 5B). With M559R-TBK1 expression, a minority of OPTN rings were positive for TBK1, phospho-OPTN, or both (Figure 5D,F). The evidence that any phospho-OPTN is present in D135N-, G217R-, and M559R-TBK1 expressing cells is surprising given the *in vitro* finding that TBK1 with these mutations cannot carry out phosphorylation (8), even while it can be recruited to ubiquitinated mitochondria (Figure 3B). While we cannot rule out contributions from residual levels of endogenous TBK1, it may also be the case that another kinase can phosphorylate OPTN in the absence of TBK1 activity.

There is a high structural similarity between the N-terminal domain of FAK family kinase-interacting protein of 200 kD (FIP200) and the scaffolding dimerization domain of TBK1. In forming the ULK1 complex, the component kinase ULK1 associates with FIP200 in an analogous position to the kinase domain of TBK1 (33). Thus, we wondered if the ULK1 complex could contribute to phosphorylation of OPTN during mitophagy. We inhibited the ULK1 complex with a specific inhibitor, ULK-101 (40) in cells depleted of TBK1 and expressing WT-TBK1, the engineered kinase-inactive mutant D135N-TBK1, or with no rescue (Figure 6A).

**Figure 6.**
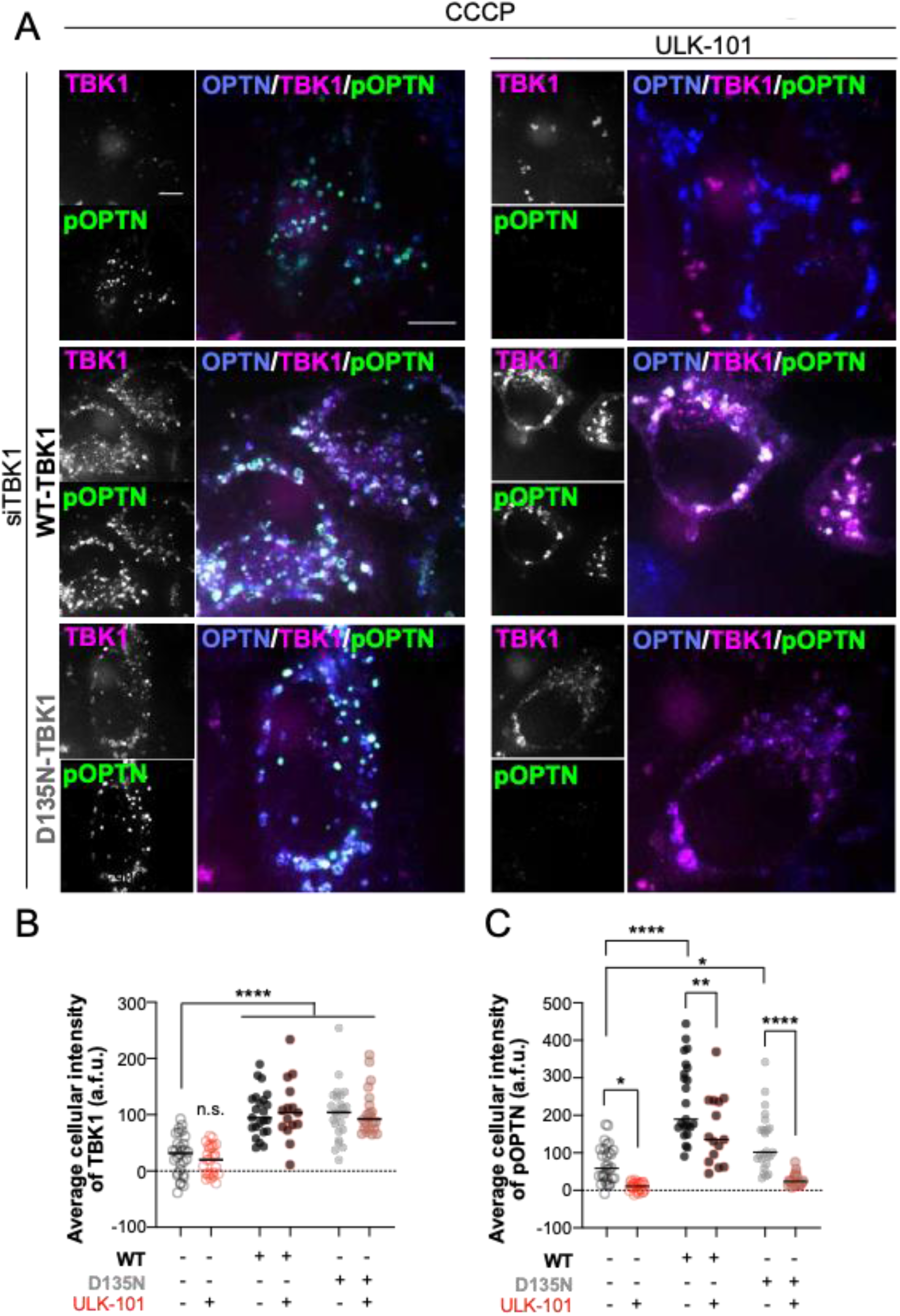
ULK1 contributes to OPTN phosphorylation independent of TBK1 kinase activity. A. Maximum intensity projection images of HeLa cells depleted of endogenous TBK1 and expressing Parkin (not tagged) and OPTN (blue). Phospho-S177-OPTN was tagged with an antibody (green). In the top row, cells were not rescued with exogenous TBK1; magenta channel shows fluorescent ligand alone. In the middle and bottom rows cells were rescued with WT- and D135N-TBK1, respectively (magenta). Half of each set was treated with the ULK1 complex inhibitor ULK-101 (right column) and all were fixed after treatment with CCCP for 90 min. Scale bars, 8 μm. B,C. Whole cell average intensities of TBK1 (B) or pOPTN (C) signal after background subtraction were measured for each condition. Bars indicate medians. n= 8-15 cells from at least 3 independent experiments. n.s., not significant (n.s. where not specified), * p ≤ 0.05, ** p < 0.01, **** p < 0.0001 by two-way ANOVA with multiple comparisons.

Corroborating our earlier results (Figure 3), WT-TBK1 and D135N-TBK1 were recruited to the same extent after mitochondrial damage, as measured by whole-cell average intensities of TBK1 (Figure 6B). Treatment with ULK-101 did not affect TBK1 recruitment. However, the persistent levels of phospho-S177 OPTN observed even in cells depleted of TBK1 (Figure 6C) was abrogated upon treatment with the ULK1 inhibitor ULK-101. As expected, cells expressing WT-TBK1 exhibited higher levels of phospho-S177 OPTN after CCCP treatment, however ULK-101 treatment diminished the average intensity of phospho-OPTN. Phospho-OPTN intensity was higher in cells expressing D135N-TBK1 compared to cells depleted of TBK1, indicating that recruitment of inactive TBK1 is sufficient to induce OPTN phosphorylation. Strikingly, inhibition of ULK1 in D135N-TBK1-expressing cells reduced phospho-OPTN to the same level seen in TBK1-depleted cells treated with the ULK1 inhibitor. These results suggest that ULK1 complex activity contributes to OPTN phosphorylation. Further, since expression of kinase-inactive D135N-TBK1 was sufficient to increase phospho-OPTN levels, we hypothesize that TBK1 recruitment facilitates ULK1 activity, leading to phosphorylation of OPTN, albeit to a lower extent. Thus, we propose that ALS-linked mutants G217R- and M559R-TBK1, diminish phospho-OPTN levels because they are not recruited to damaged mitochondria.

### TBK1 and phospho-OPTN are both required for efficient LC3 recruitment

The sequence of molecular events investigated thus far serve to induce formation of a double membraned autophagosome classically identified by membrane-associated LC3 (41). After sufficient expansion, the membrane engulfs the damaged organelle (Supplemental Figure 1D) before fusing with acidic lysosomes to complete the degradation process. In order to test whether TBK1 mutant variants would alter autophagosome engulfment, we quantified the percentage of LC3-positive mitochondria after CCCP treatment. With WT-TBK1 expression, clear recruitment of LC3 was observed to 10 ± 0.9% of damaged mitochondria in a single confocal section at the 90 min time point, and could be visualized as rings coincident with TBK1 (Figure 7A-C, Supplemental Figure 7). Expression of the monomeric R357Q-TBK1 did not result in a statistically significant change in percent of mitochondria engulfed in LC3 rings (Figure 7B).

**Figure 7.**
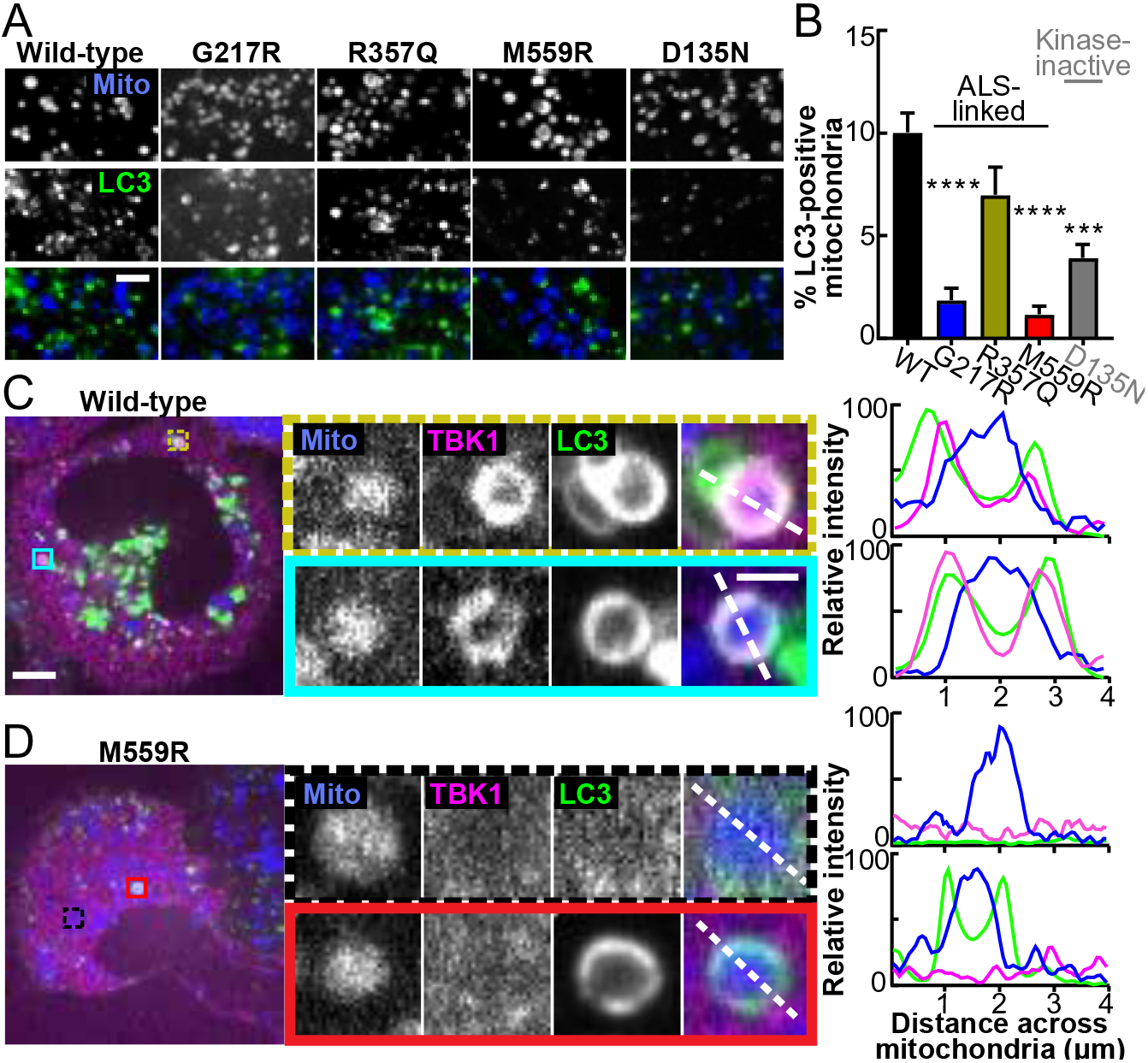
TBK1 recruitment and phosphorylation of OPTN are both necessary for efficient mitochondrial engulfment by the LC3-positive autophagosome. A. Representative confocal images of fixed HeLa cells depleted of endogenous TBK1 and expressing Parkin (not-tagged), a fluorescent mitochondrial marker (blue), LC3 (green), and TBK1 (respectively indicated above each column, channel not shown), fixed after treatment with CCCP for 90 min. Scale bar, 3 μm. B. Percent of LC3-positive mitochondria in cells expressing the respective TBK1 mutants. n= 5-15 cells from at least 3 independent experiments. *** p < 0.001, **** p < 0.0001 by ordinary one-way ANOVA with Dunnett’s multiple comparisons test. Error bars indicate SEM. C-D. Confocal images of fixed HeLa cells in our mito-damage paradigm expressing Parkin (not-tagged), a fluorescent mitochondrial marker (blue), WT- (C) or M559R- (D) TBK1 (magenta), and LC3 (green). Insets (colored boxes and zoom panels) display examples of mitophagy events. The adjacent traces (right) display quantification of relative signal intensity of each channel over a line scan (white dashed lines) across the diameter of the rounded mitochondria. Scale bars, zoom out, 10 μm; zoom in, 2 μm. Images shown in (A), (C), and (D) are insets; for representative images of whole fields, see Supplemental Figure 7.

In contrast, expression of G217R-TBK1 and M559R-TBK1 mutants significantly inhibited the formation of LC3 rings on damaged mitochondria, with 1.9% ± 0.6 and 1.2% ± 0.4 LC3-positive mitochondria, respectively (Figure 7A,B). Interestingly, while no M559R-TBK1 rings were detectable, some mitochondria in M559R-TBK1-expressing cells were LC3-positive (Figure 7D, bottom inset). The formation of LC3-positive mitochondria even in the absence of TBK1 can potentially be explained by a compensatory mechanism in which a different mitophagy receptor, NDP52 is recruited to ubiquitinated mitochondria independently of OPTN and TBK1, and can recruit LC3 by an alternative mechanism (10). However, this alternative mechanism is less efficient, as there were many fewer LC3-positive mitochondria.

Expression of the kinase-inactive D135N-TBK1 construct also impaired LC3 recruitment to damaged mitochondria. There were significantly fewer LC3-positive mitochondria with D135N-TBK1 rescue (3.9% ± 0.65) compared to WT-TBK1 (Figure 7B). The residual autophagosome formation observed in cells expressing the kinase-inactive variant is consistent with our data that recruitment of inactive TBK1 facilitates partial, ULK1-dependent OPTN phosphorylation. We conclude that TBK1 recruitment to damaged mitochondria is required, and that one of the key targets of TBK1, OPTN, must be phosphorylated in order to activate autophagosomal formation in an efficient manner.

### Expression of ALS-associated TBK1 mutants alters mitochondrial network health and sensitivity to oxidative stress

Given that TBK1 missense mutations differentially affected mitophagy when expressed in HeLa-M cells, we asked how expression of these mutants would affect mitochondrial homeostasis in primary neurons. We depleted endogenous TBK1 from primary rat hippocampal neurons and expressed constructs encoding WT-, R357Q-, or M559R-TBK1 (Supplemental Figure 8A-C). While the R357Q- and M559R-TBK1 constructs were expressed in a lower percentage of neurons than WT-TBK1, expression within individual neurons was similar for all constructs examined as quantified by cellular fluorescence intensity measurements (Supplemental Figure 8D).

First, we examined mitochondrial network health under basal condtions, focusing on somal mitochondria as our previous work has shown most mitophagic events occur within this compartment (30). We assessed network health using the polarization-dependent mitochondrial dye, tetramethylrhodamine ethyl ester (TMRE) (Figure 8A, top panels). TMRE intensities for neurons expressing R357Q-TBK1 were significantly reduced compared to WT-TBK1 under basal conditions, indicating that mutant expression is sufficient to negatively impact network health (Figure 8A,B). Of note, there was no correlation between levels of TBK1 expression and TMRE intensity at the cellular level (Supplemental Figure 8D), and no loss of somal mitochondrial mass in neurons expressing any of the variants under these conditions (Figure 8C).

**Figure 8.**
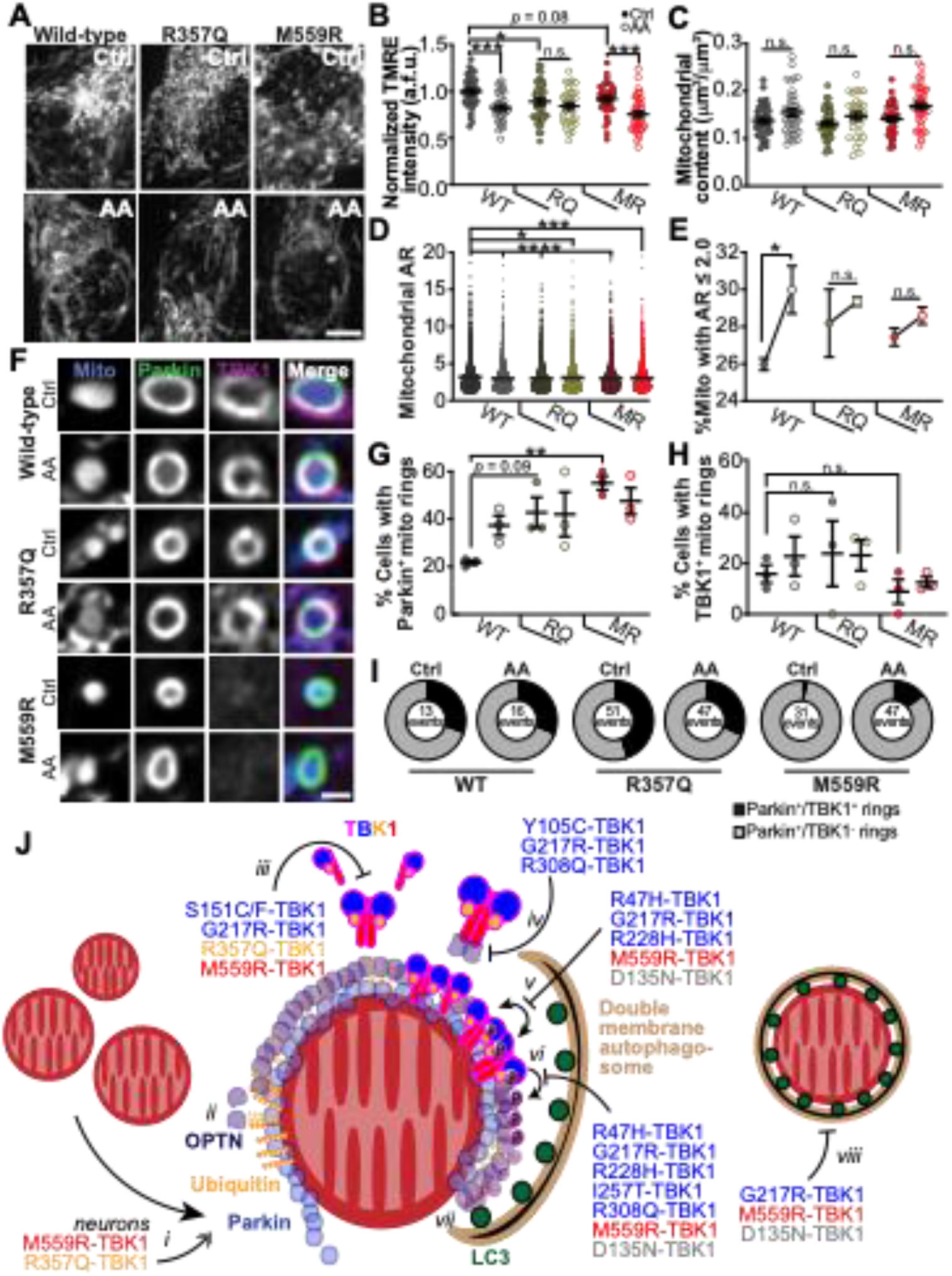
Expression of ALS-associated TBK1 mutants alters mitochondrial network health and sensitivity to oxidative stress, and model for the deleterious effects of TBK1 mutations in mitophagy. A,B. Representative images (A) and quantification (B) of TMRE fluorescence intensity. Mean ± SEM; n= 30-42 neurons from 3-4 biological replicates; 7 DIV. Not significant (n.s.), * p ≤ 0.05, *** p < 0.001 by one-way ANOVA with Sidak’s multiple comparisons test. Scale bar, 5 μm. C. Quantification of the mitochondrial content. Mean ± SEM; n= 30-42 neurons from 3-4 biological replicates; 7 DIV. Not significant (n.s.) by Kruskal-Wallis ANOVA with Dunn’s multiple comparisons test. D. Quantification of the mitochondrial aspect ratio (AR) for all mitochondria observed. Mean ± SEM; n =30-42 neurons from 3-4 biological replicates; 7 DIV. * p ≤ 0.05, *** p < 0.001, **** p < 0.0001 by one-way ANOVA with Dunnett’s multiple comparisons test. E. Percent of mitochondria with a mitochondrial aspect ratio (AR) ≤2. Mean ± SEM; n =30-42 neurons from 3-4 biological replicates; 7 DIV. Not significant (n.s.); * p ≤ 0.05 by one-way ANOVA with Sidak’s multiple comparisons test. F. Representative images of Parkin-positive mitochondria with examples that are TBK1-positive (WT and R357Q) and TBK1-negative (M559R). Scale bar, 1 μm. G,H. Quantification of the percent of neurons with Parkin-positive (G) or TBK1-positive (H) mitochondria rings. Mean ± SEM; n =25-32 neurons from 3 biological replicates; 7 DIV. Not significant (n.s.); ** *p* < 0.01 by one-way ANOVA with Sidak’s multiple comparisons test. I. Quantification of the number of Parkin-positive mitochondria (total) that are TBK1-positive (black sector) or TBK1-negative (grey sector). n =25-32 neurons from 3 biological replicates; total number of events are shown; 7 DIV. J. Model for TBK1 involvement in mitophagy, and effects of mutants. *i*) Upon mitochondrial depolarization, Parkin (blue circles) stabilizes on the outer mitochondrial membrane and proceeds to ubiquitinate (gold spheres) outer membrane proteins. In neurons, expression of R357Q- or M559R-TBK1 induces more rounded, Parkin-positive mitochondria. *ii*) Ubiquitin chains recruit OPTN (purple circles), which interact with ubiquitin via their UBAN domains. TBK1 is not required for this interaction. *iii*) TBK1 (multi-colored cartoon) monomers constitutively dimerize along their SDD domains. Five mutations disrupt this dimerization, including R357Q-TBK1 and M559R-TBK1, which have completely disrupted dimerization. *iv*) TBK1 associates with OPTN at its CTD, thus TBK1 may be co-recruited with OPTN to the mitophagy site. Three ALS-linked mutants of TBK1 have disrupted OPTN association, yet Y105C-TBK1 and R308Q-TBK1 can still be recruited to the damaged mitochondria, thus TBK1 can also be independently recruited. *v*) Formation of TBK1 multimers at the mitochondria surface promotes TBK1 trans-autophosphorylation, by which TBK1 is activated upon phosphorylation at S172 (purple circles with “P”). Four ALS-linked TBK1 mutants and the engineered D135N-TBK1 have diminished or abolished activation. *vi*) Activated TBK1 phosphorylates the mitophagy receptor, OPTN at S177. Activated TBK1 may also play a role in promoting autophagosomal membrane expansion (tan crescent). vii) Phosphorylated OPTN is necessary to recruit the LC3-coated (dark green circles) autophagosome. viii) The double membrane autophagosome completely engulfs a damaged mitochondria.

We then induced mitophagy in hippocampal neurons by applying 3 nM Antimycin A (AA) over 2 hrs (Figure 8A, bottom panels), as performed previously (30). Following treatment with AA, neurons expressing either WT-or M559R-TBK1 exhibited significantly lower intensities of TMRE (Figure 8A,B), than were observed under basal conditions. The TMRE intensity of the somal mitochondrial network in R357Q-TBK1 expressing neurons did not significantly decrease with AA treatment (Figure 8B). As an additional measure of mitochondrial damage, we analyzed mitochondrial morphology by measuring the aspect ratios (AR) of mitochondria in neurons expressing WT-, R357Q-, or M559R-TBK1 under control conditions or when treated with AA. All conditions exhibited significantly decreased ARs as compared to mitochondria in neurons expressing WT-TBK1 under basal conditions (Figure 8D), evidence that either mitochondrial depolarization with AA or expression of mutant TBK1 is sufficient to alter mitochondrial network properties. As mitochondrial rounding is a measure of stress, we focused specifically on the percentage of mitochondria with an AR of ≤2. Mutant-expressing cells tended to exhibit more rounded mitochondria than WT expressing cells in basal conditions, corroborating our observation that mutant-expressing neurons exhibit mitochondrial stress (Figure 8B,E). Given that mitochondria were already more rounded with expression of mutant TBK1 under control conditions, only neurons expressing WT-TBK1 exhibited a significant increase in rounding after AA-induced mitochondrial depolarization (Figure 8E).

Depolarized mitochondria effectively recruited both Parkin and TBK1 in neurons expressing WT-TBK1 (Figure 8F) as expected (30). In striking contrast, expression of either R357Q- or M559R-TBK1 was sufficient to cause increased Parkin recruitment to mitochondria even under basal conditions, while mitochondrial depolarization by AA did not further increase the number of Parkin rings associated with somal mitochondria (Figure 8F,G). Since transient Parkin expression may upregulate mitophagy and affect TBK1 recruitment in neurons, we identified cells with fewer than ten Parkin-positive mitochondria per soma to measure whether there was TBK1 recruitment to these events. 20-30% of neurons expressing WT- and R357Q-TBK1 exhibited mitochondria that recruited TBK1 (Figure 8H). Despite 50-60% of M559R-TBK1 expressing neurons exhibiting Parkin-positive rings (Figure 8G), only 10-15% of cells had TBK1-positive rings (Figure 8H). We looked more closely at individual events, in which depolarized, rounded mitochondria recruited Parkin (Figure 8F,I). R357Q-TBK1, like WT-TBK1, was recruited to one third of all Parkin-positive mitochondria, while M559R-TBK1 was recruited to less than a tenth of Parkin-positive mitochondria (Figure 8I). R357Q-TBK1 may inhibit mitochondrial clearance if its monomeric property induces less efficient interactions with the depolarized mitochondria even though it is recruited to the same proportion of events as WT-TBK1. This would result in more cells with Parkin-positive, TBK1-positive mitochondria, as observed (Figure 8I). In contrast, M559R-TBK1 expression results in loss of TBK1 recruitment, (Figure 8F,H,I) which may result in a more severe mitophagy defect.

## DISCUSSION

Here we present a functional analysis of recently identified and characterized ALS- and ALS-FTD-linked TBK1 mutations (8). Intriguingly, these mutations are located throughout the structure of the TBK1 molecule and result in diverse biochemical consequences, with differential effects on dimerization, auto-phosphorylation, OPTN association, and kinase activity. While TBK1 mutations may contribute to neurodegenerative pathogenesis through a number of different pathways, our study is the first to compare the effects of many of these mutations on the clearance of damaged mitochondria via mitophagy (Figure 8I), a process thought to be crucial to maintaining neuron health. We build upon previous work to propose a model of TBK1 activity in mitophagy that reinforces the hypothesis that disordered mitochondrial clearance plays a role in the development of ALS.

Previous work has suggested that TBK1 dimerization promotes kinase activation (13–15). However, most of the TBK1 mutants known to disrupt dimerization were recruited to damaged mitochondria and formed rings with the same robustness as WT-TBK1. Notably, the ubiquitin-like domain mutant R357Q-TBK1 was shown to be fully monomeric (8), yet R357Q-TBK1 rings formed with the same prevalence as WT-TBK1 rings in HeLa-M cells, and R357Q-TBK1 was recruited to the same proportion of Parkin-positive mitochondria in hippocampal neurons. In hippocampal neurons, however, expression of R357Q-TBK1 was sufficient to induce mitochondrial stress and fragmentation in both basal and oxidative stress conditions. Thus, even as the effects of inhibited dimerization on mitophagy are subtle in HeLa cells, they become magnified in more specialized cell types.

OPTN depletion prevents TBK1 recruitment to damaged mitochondria and inhibits efficient autophagosome formation (10). However, the the TBK1 mutations Y105C and R308Q reduce association of TBK1 with OPTN (8), yet both mutants were recruited to damaged mitochondria. Thus, TBK1 association with its adaptors may be more complex than previously thought. TBK1 interacts in a mutually exclusive manner with OPTN or another adaptor at its CTD, however, the C-terminal TBK1 mutation E696K affects only OPTN association and not NAP1 association (7, 16). Future experiments should investigate whether ALS-TBK1 mutations predispose TBK1 to associate with NAP1, Sintbad, or TANK instead of OPTN, or vice versa, and how this balance could impact the functional roles of TBK1.

TBK1 recruitment is thought to be necessary for OPTN phosphorylation, which enhances the affinity of OPTN for ubiquitin chains (7, 18) and is required for its interaction with LC3 (10, 18, 27). We found that partial phosphorylation of OPTN was observed even with expression of mutants that abolished TBK1 recruitment. We used the ULK1 inhibitor ULK-101 to demonstrate that this limited OPTN phosphorylation is dependent on ULK1 complex activity, suggesting that the ULK1 kinase may directly phosphorylate OPTN. This novel finding suggests that the kinase-dependent regulation of mitophagy is not simply a linear pathway, but instead the activities of TBK1 and ULK1 are inter-dependent to some degree, a possibility that will require further work to explore.

Generation, recruitment, and engulfment by the LC3-marked phagophore is the final step before the new mitophagosome compartment fuses with acidic lysosomes. Expression of the R357Q-TBK1 mutation, which is recruited to damaged mitochondria and has a functional kinase, despite its monomeric form, promotes LC3 recruitment to damaged mitochondria at WT levels in HeLa-M cells. Expression of the ALS-associated G217R and M559R mutations leads to significantly fewer LC3-positive damaged mitochondria after global oxidative damage. However, our data show that phospho-OPTN is associated with damaged mitochondria in G217R- and M559R-TBK1 expressing cells with the same prevalence as is found in WT-TBK1 expressing cells.Thus, our findings highlight the requirement for both TBK1 kinase activity and TBK1 recruitment in order to promote autophagosomal engulfment of damaged mitochondria.

TBK1 mutants that did not measurably affect mitophagy in these experiments may induce more subtle defects that only emerge over a longer period of time or in specialized cells, as we saw with the differing effects of R357Q-TBK1 expression between HeLa-M cells and hippocampal neurons. This disparity could be due to uniquely sensitive roles of the protein in different cell types; alternatively, some of the missense mutations in TBK1 may induce misfolding or protein instability, and thus decreased expression levels. Previous work has shown that some heterozygous mutations in TBK1 that produce premature stop codons, frameshifts, or in-frame deletions cause ALS by haploinsufficiency (45). We performed imaging analyses on cells with similar expression levels of tagged TBK1 constructs (Supplemental Figure 2A,B; Supplemental Figure 8D), however we did note that cell lysate analyses revealed some of the mutants examined were poorly expressed compared to WT-TBK1 (Supplemental Figure 2C; Supplemental Figure 8C). While in HeLa-M cells, some inefficiencies might be compensated by a high concentration of mutant TBK1, in neurons, a lower availability of the monomeric R357Q-TBK1 could be unable to form the proper scaffold needed for ULK1 complex regulation. This would lead to deficient phosphorylation of OPTN and inadequate recruitment of the autophagosome, despite the equivalent kinase activity of R357Q to wild-type TBK1. It will be important to examine endogenous expression levels of TBK1 in patient-derived material to more accurately distinguish between loss-of-function and haploinsufficiency.

Finally, though our study focuses on TBK1 recruitment kinetics and phosphorylation of OPTN, recent work points to other roles for TBK1 within mitophagy. TBK1 also phosphorylates RAB7A to recruit ATG9-positive vesicles as a source of autophagosomal membrane (42), facilitates the interaction of NDP52 with the FIP200/ULK1 complex to promote ULK1 activation (43) and phosphorylates LC3C and GABARAP-L2 to ensure a steady availability of the autophagosome membrane (44), each of which may also be affected by TBK1 mutations. It is also possible that mutations in TBK1 disrupt other critical pathways, such as inflammation. In the NF-κB response pathway, TBK1 is required to interact with TNF receptor-associated factor 2 (TRAF2) and TANK (46); this network may be hindered by missense mutations in TBK1. Inflammatory and viral response mechanisms may intersect or converge with mitophagy, or be wholely separate from the pathway of mitochondrial clearance, leading to varying presentations of the same disease (47, 48). Interestingly, two independent patients with the same TBK1 mutation presented with similar phenotypes of ALS (3), pointing toward the merit of further systematic correlation of genetic mutations with disease presentations.

Overall, our results indicate that some TBK1 mutations disrupt mitophagic flux, inhibiting or delaying clearance of the damaged organelles. We also noted that TBK1 mutant expression in primary neurons was sufficient to induce stress within the mitochondrial network. The accumulation of dysfunctional mitochondria may deplete cellular energy pools and/or produce cytotoxic ROS, triggering neurodegeneration. It is also possible that a disruption in flux could lead to sequestration of OPTN, TBK1, or other mitophagy components on damaged mitochondria, preventing these proteins from carrying out other cellular roles (49, 50). Deficient mitophagy may also stimulate innate immune pathways (51, 52) and promote build-up of toxic aggregates (53), provoking neuro-inflammation, another hallmark of ALS (54). Critical questions persist regarding how dysfunctional mitochondria, neuroinflammation, and toxic aggregates relate to one another in the pathogenesis of ALS. Moreover, the relative roles of neural and glial dysfunction, age of onset, and exacerbating factors such as a ‘second hit’ (55, 56) must be explored. Further dissection of those phenomena will orient our approach to therapeutic development in the future.

## ACKNOWLEDGEMENTS

We would like to thank Dr. Mariko Tokito for re-engineering the TBK1 constructs with various tags, and the entire Holzbaur group for invaluable discussion. We gratefully acknowledge Project ALS for initiating this collaboration and Project ALS (Grant ID 2018-03) and NINDS (NS060698) for supporting this work. The study is also funded by the joint efforts of The Michael J. Fox Foundation for Parkinson’s Research (MJFF) and the Aligning Science Across Parkinson’s (ASAP) initiative. MJFF administers the grant ASAP-000350 on behalf of ASAP and itself. C.S.E. was supported by the Howard Hughes Medical Institute Hanna H. Gray Fellowship.

## MATERIALS AND METHODS

Mutated TBK1 constructs were generated as described in Ye et al. (8). SNAP-tagged and Halo-tagged versions of TBK1 constructs were generated by inserting SNAP (pSNAPf [New England Biolabs]) or (Halo [pHaloTag vector, Promega]) at the N-terminus of the TBK1 coding region. HeLa-M or HEK293T cells were transfected with exogenous constructs 24 hours before sample collection. Hippocampal neurons were transfected 36-48 hours before imaging and collection. Mitochondrial enrichment was performed with ThermoScientific isolation kit for cultured cells (89874). HeLa-M cells and neurons were labeled with fluorescent ligands prior to treatment. Where applicable, fixation was done with 4% paraformaldehyde after CCCP treatment. Confocal microscopy was performed on an UltraView Vox spinning disk confocal system and images were deconvolved with Huygens Professional Software, then analyzed with ImageJ/FIJI and Ilastik software. Notably, intensity measurements were collected from original data, not deconvolved images. For details regarding all materials and methods, see extended section.

## Supplementary Information for

### MATERIALS AND METHODS (EXTENDED)

#### Reagents

Constructs used were: Mito-DsRed2 (kindly provided by. T. Schwartz, Harvard Medical School, Boston) and SBFP2-mito (Mito-DsRed2 recloned into pSBFP2-C1, Addgene #22880); Mito-SNAP (recloned from Mito-DsRed into a pSNAPf [New England Biolabs]), YFP-Parkin (a gift from R. Youle, NIH, Bethesda, MD) and untagged Parkin (subcloned from YFP-Parkin); pEGFP-OPTN (kindly provided from I. Dikic, Goethe University, Frankfurt), Halo-OPTN (subcloned from EGFP-OPTN to a pHaloTag vector, Promega); pEGFP-LC3B (a gift from T. Yoshimori, Osaka University, Osaka); and SNAP-tagged or Halo-tagged versions of all TBK1 variants. siRNA targeting the 5’ (UAACAAGAGGAUUGCCUGA) and 3’ (CCACUGUUAUACUGGGAUA) ends of hTBK1 and a scrambled control siRNA were from Horizon Discovery, and used on HeLa-M cells. ON-TARGET*Plus* Rat TBK1 (299827) siRNA *SMARTpoo*l (L-101406-02-0005; Horizon) were used on neurons. SNAP ligands (SNAP-Cell 647-SiR, S9102S and SNAP-Cell 430, S9109S) were from New England BioLabs. Halo ligands (JaneliaFluor 646, GA112A and TMR, G8252) were from Promega. TMRE (tetramethylrhodamine ethyl ester, Ethyl Ester, Perchlorate) was purchased from Life Technologies, (T-669). Antibodies used were: anti-TBK1 (abcam, ab40676, IF: 1:100, and Novus Biologicals, 108A429, WB: 1:1000), anti-SNAP (New England BioLabs, P9310, WB: 1:1000), and anti-phospho-S177-OPTN (Cell Signaling Technology, 57548, IF: 1:200). The drug carbonyl cyanide 3-chlorophenylhydrazone (CCCP) was purchased from Millipore Sigma (C2759). Antimycin A (A8674) and Oligomycin A (75351) were purchased from Sigma-Aldrich. ULK-101 (S8793) was purchased from Selleckchem.

#### Cell culture and transfection

HeLa-M cells and HEK293T cells were maintained in DMEM (Corning, 10-017-CV) with 10% fetal bovine serum (HyClone) and 1% GlutaMAX glucose supplement (Gibco, 35050061). Cells were maintained in an environment of 37 °C with 5% CO_2_. 48 hours prior to fixation or live imaging, 0.28 million cells were plated on each glass-bottomed 35 mm dish (MatTek, P35G-1.5-20-C). HEK cells were plated onto glass coverslips that had been pre-coated with poly-L-lysine for 24 hr in order to prevent sloughing. 18-20 hours prior to fixation or live imaging, cells were approximately 80-90% confluent, and were transfected with the appropriate constructs using Lipofectamine 2000 (ThermoFisher Scientific, 11668027) at a 1:4 ratio of mass (ug) to volume (uL). siRNA was transfected simultaneously at 40 μM.

#### Primary hippocampal culture and transfection

A suspension of embryonic day 18 Sprague Dawley rat hippocampal neurons were provided from the Neurons R Us Culture Service Center at the University of Pennsylvania. Cells were plated on 35 mm glass-bottom dishes (MatTek) at a density of 250,000 cells/dish; dishes were precoated with 0.5 mg/ml poly-L-lysine (Sigma Aldrich). Cells were initially plated in MEM supplemented with 10% horse serum, 33 mM D-glucose, and 1 mM sodium pyruvate and left for 2-5 hours. The media was then replaced with Neurobasal (Gibco) supplemented with 33 mM D-glucose, 2 mM GlutaMAX (Invitrogen), 100 units/ml penicillin, 100 μg/ml streptomycin, and 2% B-27 (ThermoFisher) (Maintenance Media; MM) and cells were maintained at 37 C in a 5% CO_2_ incubator. AraC (5 μM) was added the day after plating to prevent glia cell proliferation. Neurons were transfected at 5 DIV with DNA (0.8-1.2 μg of total plasmid) and siRNA (45 pmol) mixtures using Lipofectamine 2000 Transfection Reagent (ThermoFisher) and incubated 36-48 hours. To induce mitophagy, media was fully replaced with MM containing 3 nM Antimycin A for 2 hours; in control conditions media was replaced with standard MM.

#### Labeling, treatment, and fixation

##### HeLa-M and HEK cells

To tag Halo-tagged or SNAP-tagged proteins, cells were incubated with the respective Halo or SNAP ligands. For Halo, cells were incubated with 190 nM Halo ligand for at least 20 min. Cells were then washed two times with conditioned media and allowed to rest for at least 20 min in conditioned media. For SNAP, cells were incubated with 1.25 μM SNAP ligand for at least 1 hour. Cells were then washed two times with conditioned media and allowed to rest for at least 30 min in conditioned media. When both SNAP and Halo were used, cells were incubated with Halo tag first, then SNAP tag, with their respective protocols. Cells were washed two more times with conditioned media. When appropriate, cells were treated with 5 μM ULK-101 for 2.5 hr. Cells were then treated with 20 μM CCCP or a combination of 10 μM Oligomycin A and 10 μM Antimycin A in conditioned media for 1.5 hours. Immediately afterward, cells were washed with warmed PBS then fixed with warmed 4% paraformaldehyde for 10-12 min. For experiments with antibody tagging, cells were permeabilized with 0.5% Triton X at room temperature for 5 min, then blocked with 3% BSA, 0.2% Triton X for one hour. Cells were incubated with primary antibodies overnight at 4 °C. Afterward, cells were washed 4x 5 min in PBS and incubated with secondary antibodies for one hour. Cells were then washed 4x 5 min in PBS and imaged. For the transfection levels test (Supplemental Figure 2A,B), after fixation cells were labeled with Hoechst 33342 for 5 min, then again washed 4x 6 min before imaging. <u>Hippocampal neurons</u>: Prior to imaging, Halo-tag (JaneliaFluor 646, 100 nM) and SNAP-tag (SNAP-Cell 430, 2 μM) ligands were added for 30 min, followed by two quick washes and a 30 min washout. Mitochondrial membrane potential was assessed by loading mitochondria with 2.5 nM TMRE for 30 min.

#### Fixed and live cell imaging

##### HeLa-M and HEK293T cells

For the transfection levels test in HeLa-M cells (Supplemental Figure 2A,B), the cells were imaged with widefield microscopy. Images were taken from three fields per dish, and the only requirement for each field was that the nuclei (Hoechst staining) had to appear healthy and regularly spaced in an area that was close to fully confluent. For all other experiments, samples were imaged with a Nikon Eclipse Ti Microscope with a 100X objective (Apochromat, 1.49-N.A. oil immersion) and an UltraView Vox spinning disk confocal system (PerkinElmer). Z-stacks at 0.15 nm/step or timelapse confocal images at 30 seconds/frame were collected with Volocity acquisition software (PerkinElmer). Fields of view were chosen to maximize the number of cells that expressed detectable components of interest. In fixed samples, z-stacks were collected through the majority of cells’ midsections.

For experiments with live cell imaging, conditioned media was replaced with Leibovitz’s L-15 Medium (Gibco, 11415064) supplemented with 10% fetal bovine serum. Cells were then rested for at least 10 min in the 37 °C imaging chamber of the microscope. For timelapse mitochondrial damage, a z position was chosen in the midsection of a healthy-appearing cell with a regularly shaped nucleus (nucleus characterized by absence of tagged TBK1). 5-10 frames were collected at basal conditions, then a volume of imaging media at least 50% of the initial volume was added, including CCCP to bring the total concentration to 20 μM as frame collection continued.

##### Hippocampal neurons

Neurons were imaged in HibernateE (Brain Bits) supplemented with 2% B27 and 33 mM D-glucose; Antimycin A was added to the imaging media for treated conditions. TMRE was added to the imaging media for TMRE experiments.

#### Mitochondrial enrichments and immunoblots

For standard cell lysis, cells were washed two times with warmed PBS, then lysed with ice cold RIPA buffer (50 mM Tris-HCl, 1 mM EDTA, 2 mM EGTA, 1% Triton X, 0.5% sodium deoxycholate, 0.1% sodium dodecyl sulfate, 150 mM NaCl) with added Halt Protease and Phosphatase Inhibitor Cocktail (ThermoFisher Scientific, 78444) and scraped into sample tubes and incubated with continuous gentle inversion at 4 °C for 20 min. Samples were then spun at 4 °C in a microcentrifuge at 17 g for 20 min, and the supernatant was transferred to a separate tube. Samples were assayed for protein concentration with Pierce BCA Protein Assay Kit (ThermoFisher Scientific, 23225). Mitochondrial enrichment was performed with ThermoScientific isolation kit for cultured cells (89874) and mitochondrial fractions were diluted in RIPA buffer with Halt Cocktail as above.

30 μg of each sample was loaded into a 10% gel (or 14% gel for TOM20 detection). After electrophoresis, protein bands were transferred to PVDF membrane (Immobilon-FL, Millipore Sigma, IPFL00010) and total protein was labeled with Revert 700 (Li-Cor, 926-11011) and imaged on an Odyssey CLx machine (Li-Cor). After clearing the total protein stain with a solution of 0.1% NaOH, 30% methanol, the membrane was then blocked with TrueBlack blocking solution (Biotium, 23013-T) for 1 hour at room temperature. Primary antibodies were incubated overnight at 4 °C. After primary antibody incubation, the membrane was washed 4x 5 min in TBS with 0.2% TWEEN-20, and secondary fluorescent antibodies (Li-Cor IRDyes, 926-68073, 926-32212) were used at 1:20,000 for 1 hour at room temperature. Finally, the membrane was washed 4x 5 min again before it was imaged again.

#### Image processing and analysis

Images were deconvolved with Huygen’s Professional version 17.10 software (Scientific Volume Imaging, The Netherlands, http://svi.nl) to remove background noise and increase resolution and signal-to noise ratio. The Classic Maximum Likelihood Estimation (CMLE) algorithm with theoretical PSF was performed for 50 iterations at most. The signal-to-noise ratio for all channels was set between 10 and 30, depending on the individual construct; all other settings were default. Images were assembled in Illustrator (Adobe). Most images shown are deconvolved, with the exceptions of widefield images (Supplemental Figure 2A), all Supplemental whole-field images, the experiment to determine ULK1 dependence (Figure 6), and neuronal mitochondrial TMRE images (Figure 8A). All intensity measurements were carried out on raw intensity images, not deconvolved images.

In Figures 1E,F, 2, 3, 5, and 6, maximal projections of 3.75 μm were generated through the central volume of the cells. For other analyses, confocal sections were used. Parkin, OPTN, TBK1, and LC3 rings were delineated by hand and measured in ImageJ (57). Patterns of OPTN and TBK1 were classified as rings if they coincided with Parkin signal around a rounded mitochondrion. At least half of the ring had to be clearly evident to be counted. Diameters and intensities were measured for all ring ROIs in each cell and averaged. Thus, each point is the average for a single cell. To quantify percentage of LC3-positive mitochondria, Ilastik software version 1.2.2 (58) was trained to categorize mitochondria using the dsRed2-mitochondria channel in the Pixel Classification module. Feature selection used color/intensity, edge, and texture up to σ = 5 pixels. Binary images were exported as .tifs using simple segmentation, and the Analyze Particles function of ImageJ was used to count mitochondria. Mitochondria were considered positive for LC3 if the fluorescent LC3 ring surrounded at least half of the organelle. Each data point represents the percentage of LC3-positive mitochondria for a single cell. For Figure 5, OPTN rings were identified as before. Then OPTN ring regions of interest (ROIs) were transferred to the TBK1 and phospho-OPTN channels of the same image, and mean fluorescence intensity was measured for each ROI on the non-deconvolved images. Intensity data points were plotted for WT-TBK1-expressing cells, and all intensities above the 25^th^ percentile were considered “positive,” while all intensities below were considered “negative” (see Supplemental Figure 6B,C). Thus, every OPTN ring could be categorized as positive or negative for phospho-OPTN and TBK1. Images were blinded before ring counting. For ULK1 experiments, outlines were drawn around whole cells and the average intensity in the region was measured in FIJI.

For timelapse imaging, events were quantified if they remained within the z frame for most of the sequence. Background was calculated by measuring mean intensity in the JF696 channel of a ROI drawn in an area of the cell with no detectable events. This mean was subtracted from the ROI for the event at each point over the timelapse. Hence, the M559R-TBK1 signal falls below zero, since there is some photo-bleaching of the signal over time.

For volume rendering, deconvolved .tif files were converted to .ims files with the Imaris File Converter and processed with Imaris software (Oxford Instruments). Normal shading mode was used to render images of the respective volumes of mitochondria and mitophagy components of representative events.

For hippocampal neurons, following image processing, protein ring formation and mitochondrial fragments were manually identified and quantified using Fiji, where only clearly defined structures were quantified. Fluorescence intensities (TMRE and Halo-TBK1) were quantified from max projections of unprocessed z-stack images in Fiji. The values of five individual areas (2.2 x 2.2 μm square) in the soma were averaged to determine the mean gray value for each cell. Mitochondrial content was determined by dividing the mitochondrial volume by the somal volume for each cell, where volume measurements were determined using the volume measurement function in Volocity Quantitation (Quorum Technologies). The mitochondrial aspect ratio was determined using the Ridge Detection Plugin with the SLOPE method for overlap resolution on single plane images. The Particle Analyzer tool with Shape Descriptors and the Aspect Ratio (AR) function was used to quantify the mitochondrial aspect ratio for individual mitochondria within the neuron. Prism (GraphPad) was used to plot all graphs and determine statistical significance. Adobe Illustrator was used to prepare all figures and images.

Transfection level images included Hoechst and JF646 channels (Supplemental Figure 2A,B). Both channels were maximally projected, and the JF696 channel was background subtracted with ImageJ’s rolling ball radius set to 25.0 pixels, with sliding paraboloid and disabled smoothing. Images were then imported to CellProfiler software (59) and nuclei were identified as primary objects; then cells were delineated by the propagation method in the JF646 channel. Thus, only cells expressing SNAP-TBK1 were identified and the mean intensities of their cytoplasmic signals were exported to Excel. These values were displayed on a histogram to demonstrate the relative frequencies of mean intensities (GraphPad).

For immunoblots, ImageStudio Software (Version 5, Li-Cor) was used to scan bands to ensure no patches were overexposed. ImageStudio was used to subtract background and quantify band intensities, which were normalized to the total protein signal for their respective lanes with Excel (Microsoft). For mitochondrial enrichment, bands were normalized to TOM20. Those values were graphed in GraphPad (Version 9, Prism).

## SUPPLEMENTAL FIGURES

**Supplemental Figure 1.**
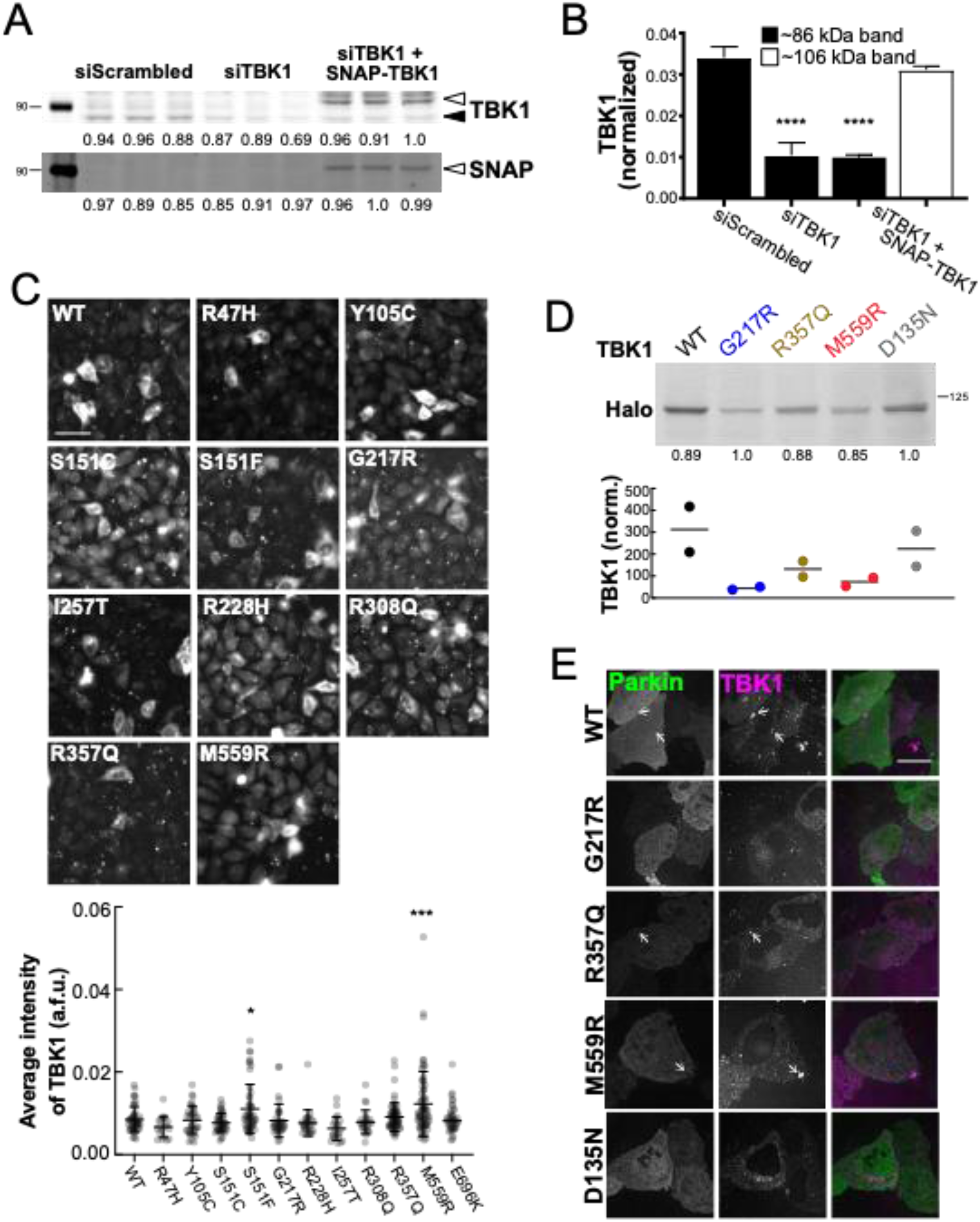
TBK1 is depleted and tagged TBK1 is exogenously expressed in cells that are induced to carry out mitophagy. A. Western blot of HeLa cells under mock, knockdown, and rescue conditions. Samples were probed for total TBK1 (top panel) and for SNAP (bottom panel). TBK1 is approximately 86 kDa (solid black arrowhead, top panel), and SNAP-TBK1 is expected to appear at 106 kDa (white arrowheads, both panels). Numbers to the left of blots indicate kDa based on protein ladder (not shown). B. Band intensities were quantified and normalized to total protein, as indicated by the numbers below each lane in (A). n= 3 independent collections. **** p < 0.0001 by ordinary one-way ANOVA with Dunnett’s multiple comparisons test. C. Representative widefield images of fixed HeLa cells in basal conditions, depleted of endogenous TBK1 and expressing siRNA-resistant SNAP-tagged TBK1 variants (grayscale). Scale bar, 60 μm. Below, graph of TBK1 average signal intensities for cells transfected with the respective TBK1 constructs. Bars indicate mean with SD. * p ≤ 0.05, *** p < 0.001 by ordinary one-way ANOVA with Dunnett’s multiple comparisons test. D. Representative Western blot of HeLa cell lysates depleted of endogenous TBK1 and expressing the respective Halo-tagged TBK1 variants (∼119 kDa). Quantification of Halo band relative to total protein shown below (n = 2). Number to the left of blot indicates kDa based on protein ladder (not shown). E. Representative confocal images of fixed HeLa cells in basal conditions, depleted of endogenous TBK1 and expressing Parkin (green) and siRNA-resistant Halo-tagged TBK1 variants (magenta). Arrows indicate TBK1 aggregates. Scale bar, 20 μm.

**Supplemental Figure 2.**
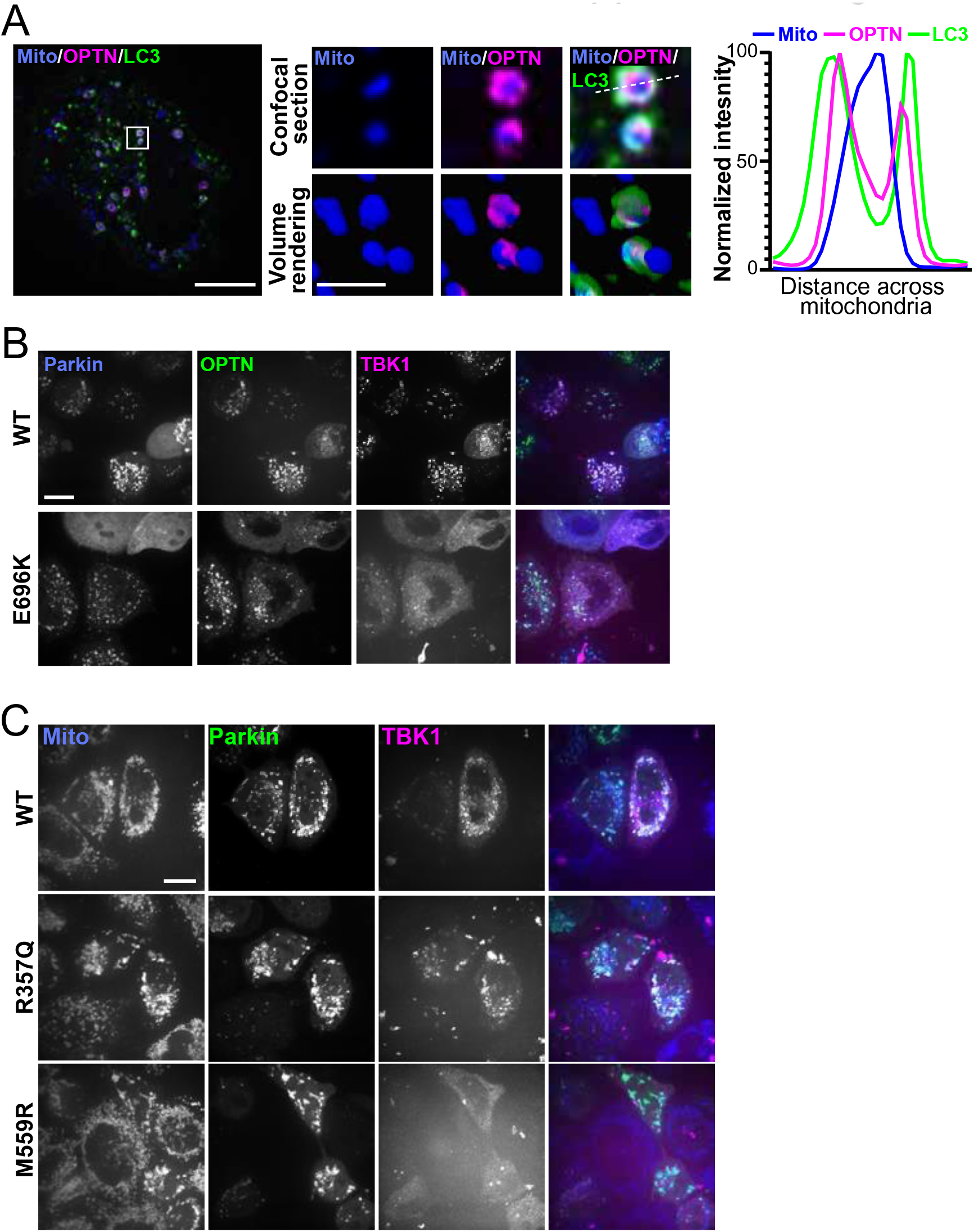
OPTN and LC3 are recruited to damaged mitochondria, and corresponding whole-field images for Main Figures 1,2. A. Confocal section of a HeLa cell expressing Parkin (not tagged), a mitochondrial maker (blue), OPTN (magenta) and LC3 (green), fixed after treatment with CCCP for 90 min. The inset (white box) and zoom images (right, top row) exhibit two mitochondria that have recruited OPTN and LC3. A volume rendering is shown below (right, bottom row). Scale bars, zoom out, 10 μm; zoom in, 2 μm. Right, profiles of relative signal intensities for mitochondria (blue line), OPTN (magenta line), and LC3 (green line) are quantified across the diameter of the rounded mitochondria (white dashed line in zoom image). B. Representative whole-field images corresponding to images in Main Figure 1E,F. Scale bar, 25 μm. C. Representative whole-field images corresponding to images in Main Figure 2A. Scale bar, 25 μm.

**Supplemental Figure 3.**
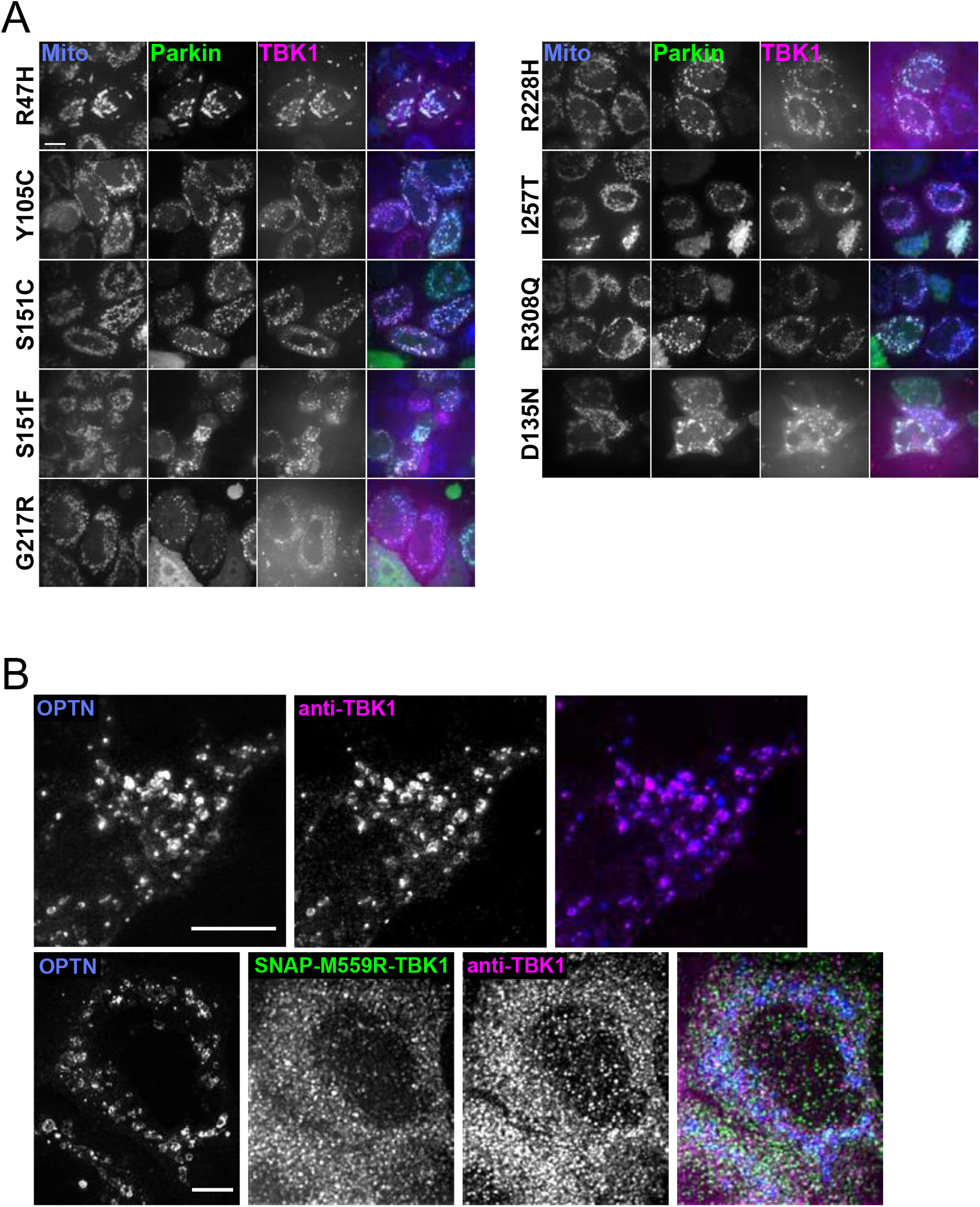
Comparison of total TBK1 recruitment to damaged mitochondria in WT-versus M559R-TBK1 expressing cells. A. Representative whole-field images corresponding to images in Main Figure 3A. Scale bar, 25 μm. B. Maximum intensity projection image of HeLa cells depleted of endogenous TBK1 and expressing Halo-OPTN (blue) and WT-TBK1 (top row, not labeled) or M559R-TBK1 (bottom row, green), fixed after 90 min treatment with CCCP. Cells were tagged with an antibody to total TBK1 (magenta). Scale bars, 10 μm.

**Supplemental Figure 4.**
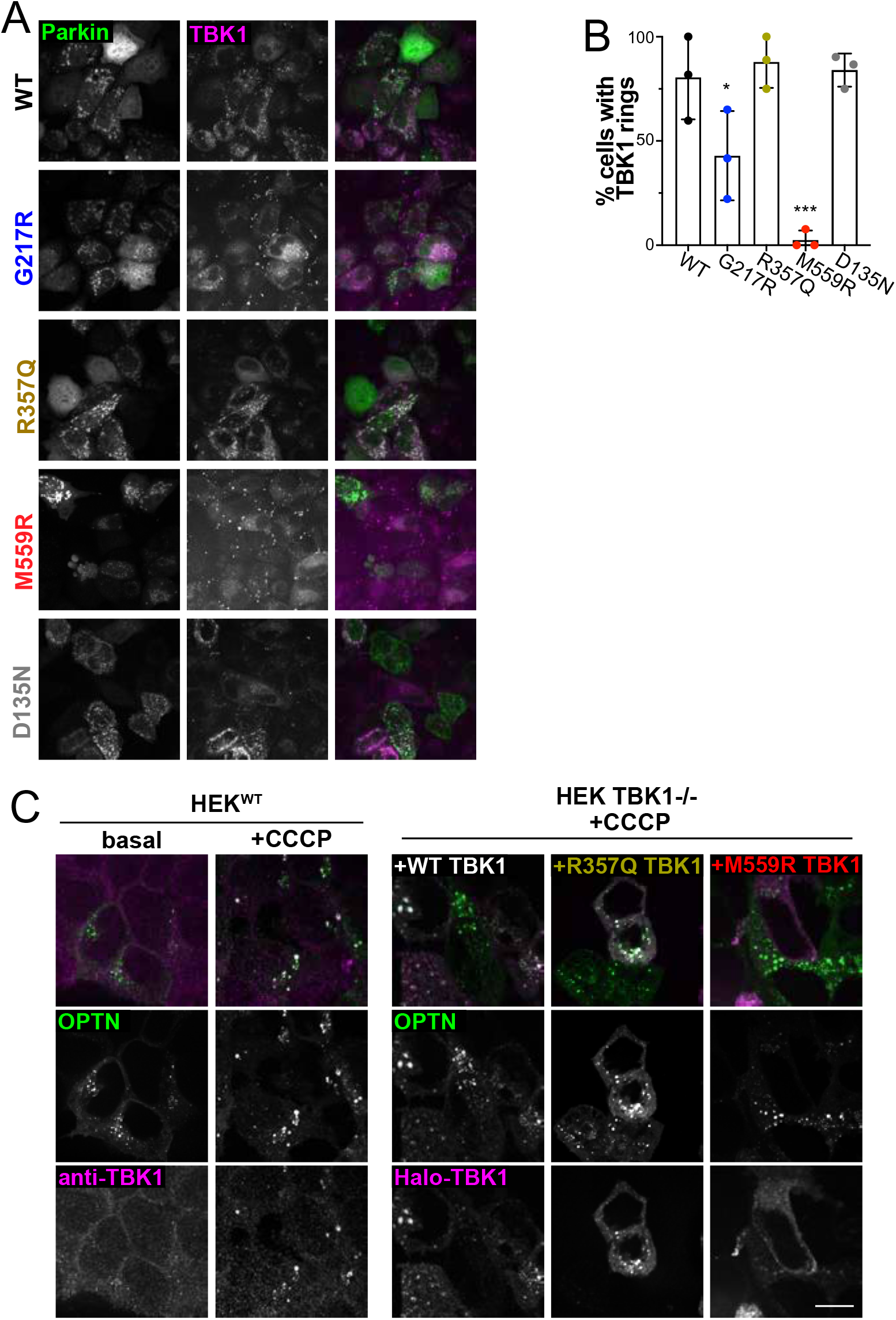
Treatment with Antimycin A and Oligomycin A induces mitochondrial depolarization and TBK1 recruitment to damaged mitochondria, and HEK cell mitochondria recruit OPTN and TBK1. A. Maximum intensity projection image of HeLa cells depleted of endogenous TBK1, expressing Parkin (green) and the respective TBK1 mutants (magenta), fixed after 90 min treatment with Antimycin A/Oligomycin A. B. Quantification of percentages of cells observed that exhibit clear TBK1 rings that coincide with Parkin. * p ≤ 0.05, *** p < 0.001 by ordinary one-way ANOVA with Dunnett’s multiple comparisons test. Error bars indicate SD. n= 3 independent experiments. C. WT or TBK1−/− HEK cells expressing EGFP-OPTN (green) and fixed. WT HEK cells were tagged with an antibody to TBK1 (magenta) in basal conditions or after 90 min CCCP treatment. TBK1−/− HEK cells expressed WT-, R357Q-, or M559R-TBK1 (magenta) before treatment with CCCP and fixation. Scale bar, 10 μm.

**Supplemental Figure 5.**
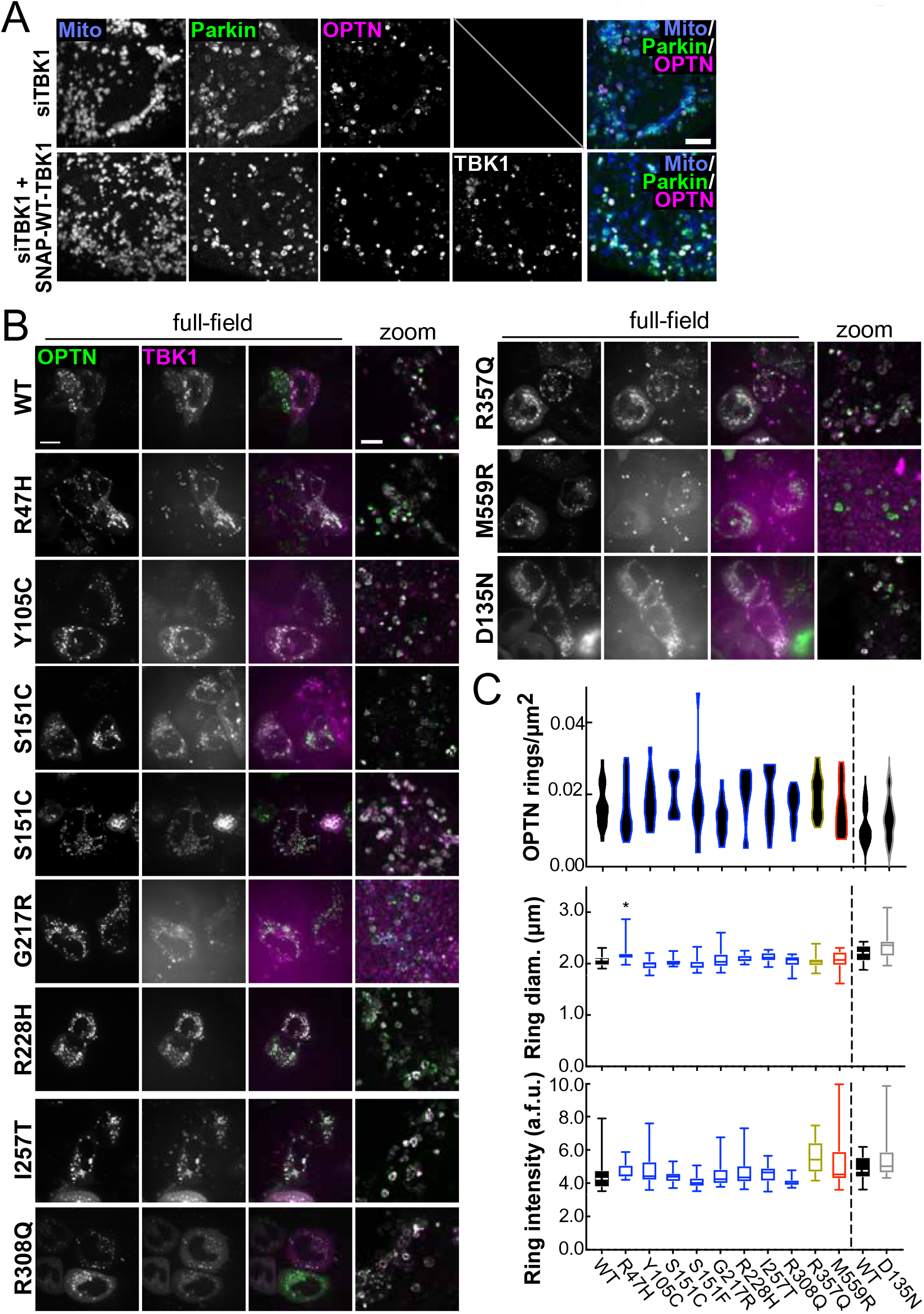
OPTN is recruited to damaged mitochondria despite depletion of endogenous TBK1. A. Maximum intensity projection images of fixed HeLa cells depleted of endogenous TBK1, expressing a mitochondrial-localizezd fluorophore (blue), Parkin (green), and OPTN (magenta), fixed after 90 min treatment with CCCP. In the bottom panels, cells were recued with exogenous SNAP-WT-TBK1, displayed in grayscale in the bottom right panel (TBK1 not included in the merged image). Scale bar, 10 μm. B. HeLa cells depleted of endogenous TBK1 and expressing Parkin (not shown), OPTN. (green), and TBK1 variants (magenta), fixed after treatment with CCCP for 90 min. First three columns are whole-field view. Scale bar, 10 μm. Final column is zoom of merged channels. Scale bar, 5 μm. C. Quantification of OPTN rings/μm^2^ (B) ring diameter (diam.) (C), and ring signal intensity (D). n= 14-20 cells from at least 3 independent experiments. * p ≤ 0.05 by ordinary one-way ANOVA with Dunnett’s multiple comparisons test. Arbitrary fluorescent units, a.f.u.

**Supplemental Figure 6.**
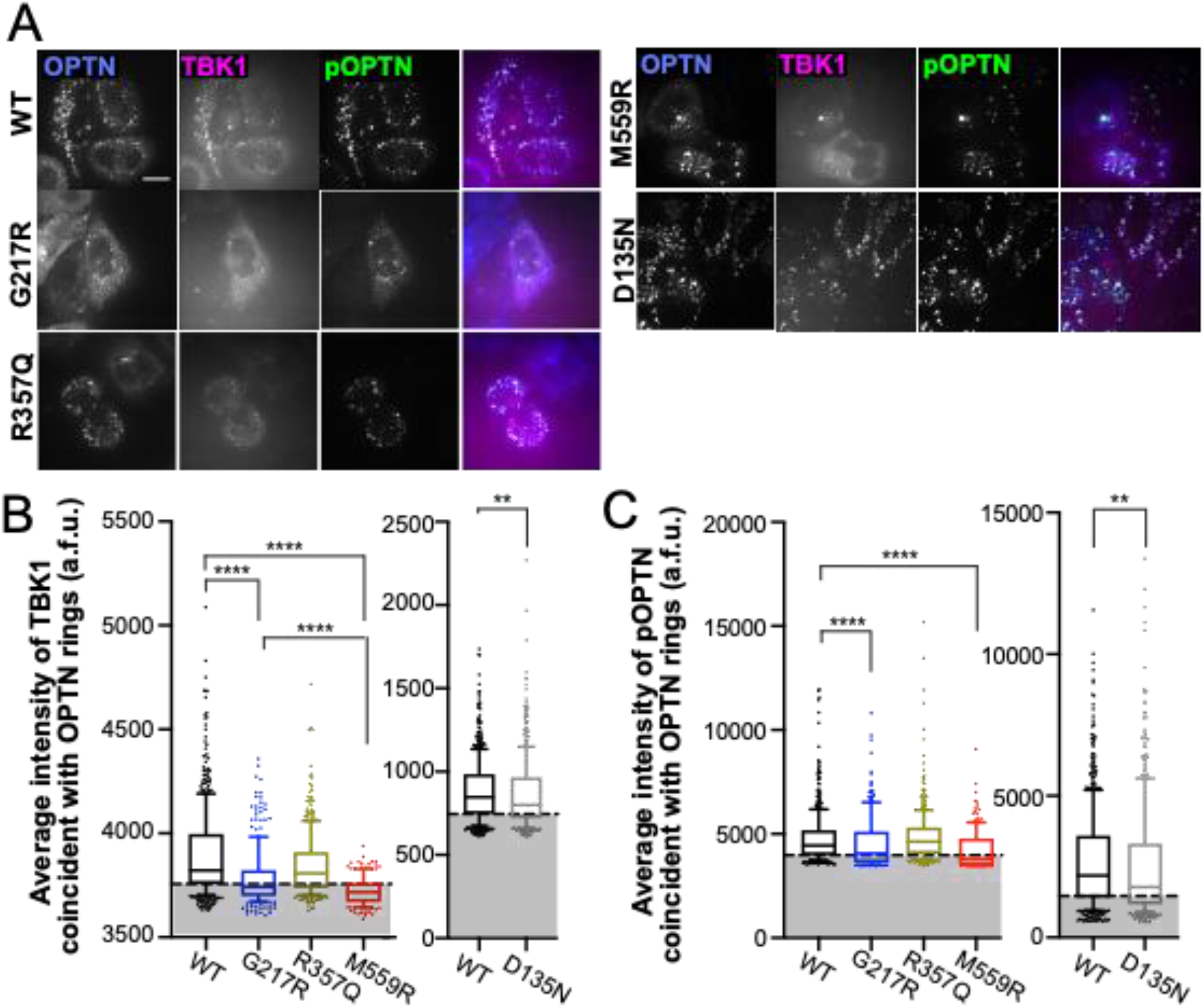
Raw intensities of TBK1 and phospho-OPTN signals with different TBK1 mutants expressed. A. Representative whole-field images corresponding to images in Main Figure 6A-E. Scale bar, 25 μm. B,C. Raw TBK1 (A) and phospho-OPTN (B) intensity measurements for OPTN rings in the respective TBK1 variant-expressing cells. Black dashed horizontal lines indicate 25^th^ percentile cutoff. Graphs are divided into experiments carried out more than one year apart. Statistical analysis among graphs of four mutant expressions (WT, G217R, R357Q, and M559R) were carried out with Kruskal-Wallis test with Dunn’s multiple comparisons. For analysis between two mutant expressions (WT and D135N), Mann-Whitney test was used. ** p < 0.001, **** p < 0.0001 Arbitrary fluorescent units, a.f.u.

**Supplemental Figure 7.**
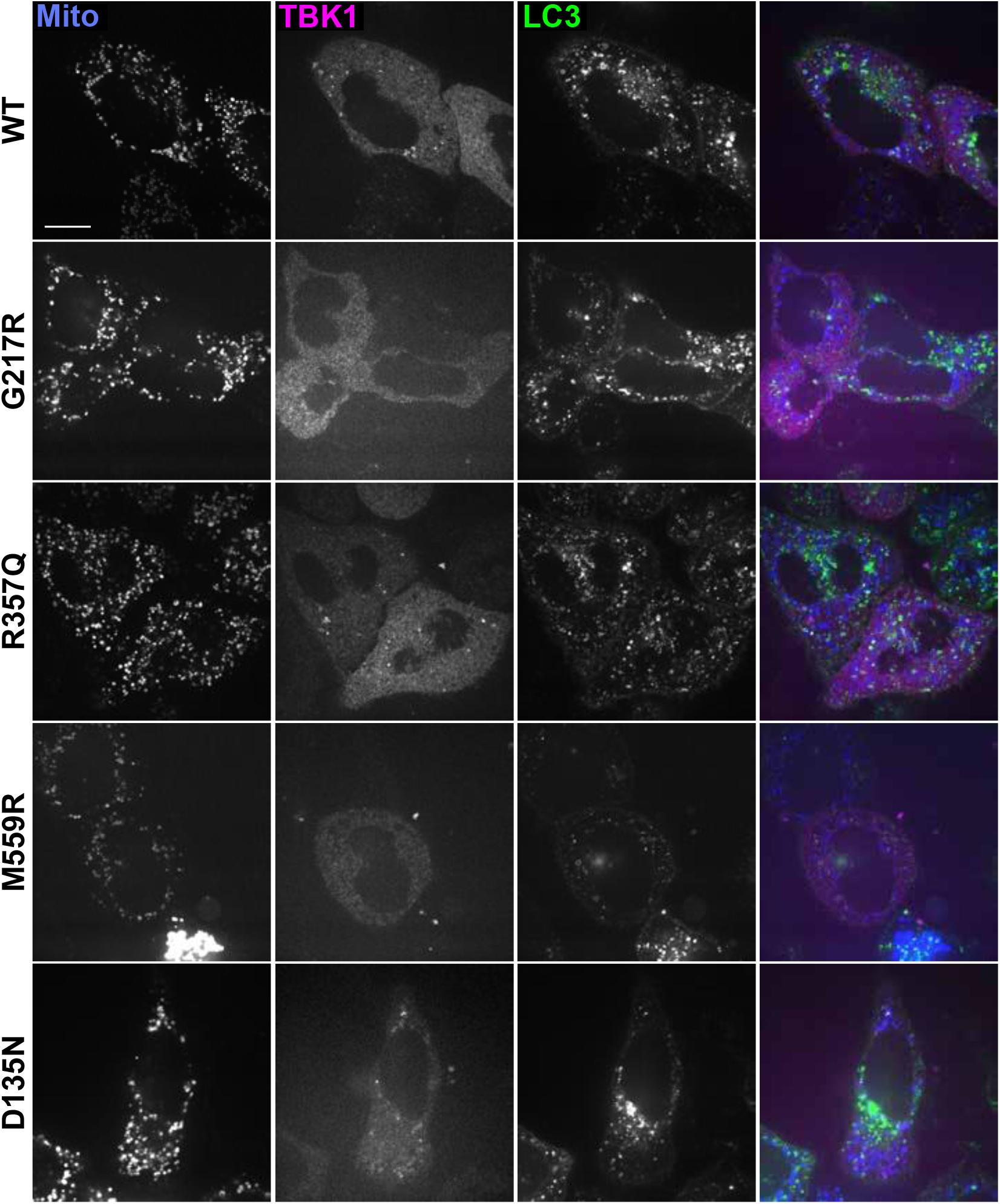
Representative whole-field images corresponding to images in Main Figure 7A,C,D. Scale bar, 25 μm.

**Supplemental Figure 8.**
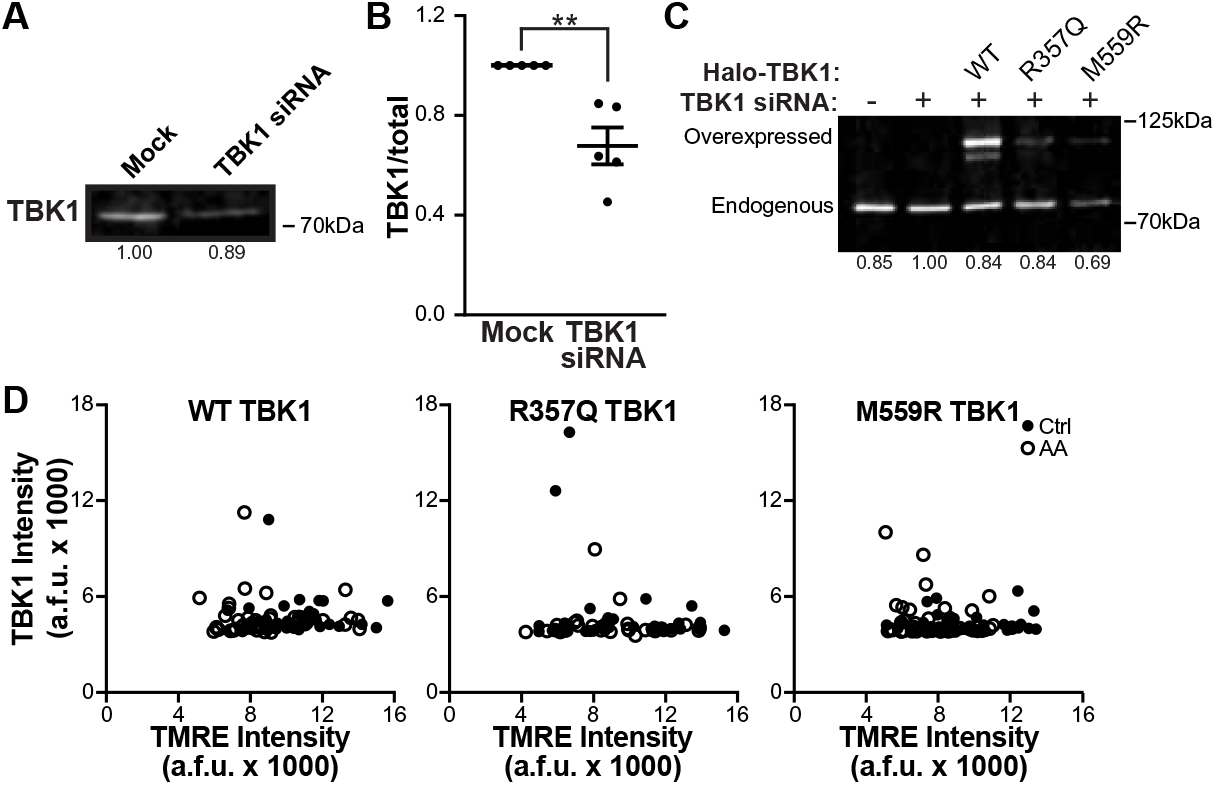
TBK1 is efficiently knocked down in neurons with siRNA. A,B. Representative Western blot (A) and quantification (B) of neurons after treatment with mock or TBK1 siRNA. Data shown as the fold change over control of TBK1 divided by total protein stain. Normalization factors are shown under lanes. Mean ± SEM; n= 5; 7 DIV. ** p < 0.01 by unpaired t test. C. Western blot of non-transfected, TBK1 siRNA treated, and TBK1 siRNA treated neurons overexpressing WT or mutant Halo-TBK1. Normalization factors are shown under lanes. D. TBK1 fluorescence intensity plotted as a function of the TMRE fluorescence intensity for each cell (data also presented Figure 8B). n= 30-42 neurons from 3-4 biological replicates; 7 DIV.

## Notes

http://doi.org/10.5281/zenodo.4670341

### Competing Interest Statement

The authors have declared no competing interest.

